# Engineering a highly selective, hemoprotein-based scavenger as a carbon monoxide poisoning antidote with no hypertensive effect

**DOI:** 10.1101/2025.01.18.633417

**Authors:** Matthew R. Dent, Anthony W. DeMartino, Qinzi Xu, Xiukai Chen, Alay Ghandi, John Hwang, Kaitlin A. Bocian, Elmira Alipour, K. Burak Ucer, Stephen R. Baker, Ajay Ram Srimath Kandada, Angka Bulbul, Daniel B. Kim-Shapiro, Jason J. Rose, Jesus Tejero, Mark T. Gladwin

**Affiliations:** Heart, Lung, Blood, and Vascular Medicine Institute, University of Pittsburgh, Pittsburgh, PA, USA; Department of Medicine, University of Maryland School of Medicine, Baltimore, MD, USA; Department of Physics, Wake Forest University, Winston-Salem, NC, USA; Translational Science Center, Wake Forest University, Winston-Salem, NC, USA; Division of Pulmonary, Allergy, and Critical Care Medicine, University of Pittsburgh, Pittsburgh, PA, USA; Department of Bioengineering, University of Pittsburgh, Pittsburgh, PA, USA; Department of Pharmacology and Chemical Biology, University of Pittsburgh, Pittsburgh, PA, USA

**Author notes:** corresponding authors: Heart, Lung, Blood and Vascular Medicine Institute, University of Pittsburgh, E1228-5A Biomedical Science Tower, 200 Lothrop Street, Pittsburgh, PA 15261, USA. E-mail address (J. Tejero), Department of Medicine, University of Maryland School of Medicine, Baltimore, MD 21201, USA. University of Maryland School of Medicine, Baltimore, MD 21201, USA. (M.T. Gladwin).

## Abstract

Carbon monoxide (CO) poisoning causes 50,000-100,000 emergency department visits and ∼1,500 deaths in the United States annually. Current treatments are limited to supplemental and/or hyperbaric oxygen to accelerate CO elimination. Even with oxygen therapy, nearly half of CO poisoning survivors suffer long-term cardiac and neurocognitive deficits related to slow CO clearance, highlighting a need for point of care antidotal therapies. Given the natural interaction between CO and ferrous heme, we hypothesized that the hemoprotein RcoM, a transcriptional regulator of microbial CO metabolism, would make an ideal platform for CO-selective scavenging from endogenous hemoproteins. We engineered an RcoM truncate (RcoM-HBD-CCC) that exhibits high CO affinity (*K*_a,CO_ = 2.8×10^10^ M^-1^), remarkable selectivity for CO over oxygen (*K*_a,O2_ = 1.4×10^5^ M^-1^; *K*_a,CO_/*K*_a,O2_ = 1.9×10^5^), thermal stability (T_m_ = 72°C), slow autoxidation rate (*k*_ox_ = 1.1 h^-1^). In a murine model of acute CO poisoning, infused RcoM-HBD-CCC accelerated CO clearance from hemoglobin in red blood cells and was rapidly excreted in urine. Moreover, infused RcoM-HBD-CCC elicited minimal hypertension in mice compared to infused hemoglobin, attributed to a comparatively limited reactivity toward nitric oxide (NO) via dioxygenation (*k*_NOD_(RcoM) = 6-8×10^6^ M^-1^s^-1^ vs *k*_NOD_(Hb) = 6-8×10^7^ M^-1^s^-1^). These data suggest that RcoM-HBD-CCC is a safe, selective, and efficacious CO scavenger. Additionally, by limiting hypertension RcoM-HBD-CCC improves end-organ adverse effects compared with hemoglobin-based therapeutics.

## INTRODUCTION

Carbon monoxide (CO) poisoning results in approximately 50,000-100,000 hospital visits in the U.S. each year, and between 1,500-2,000 people die from CO exposure annually (1, 2). Even with oxygen supplementation, the current standard of care treatment for CO poisoning, survivors often suffer increased long-term mortality (3), as well as long-term cardiac and neurocognitive deficits (4–10). The toxicity of CO is primarily attributed to a shutdown of aerobic respiration due to decreased oxygen (O_2_) delivery and utilization, combined with direct inhibition of cytochrome *c* oxidase (Complex IV) (1, 2). CO binds the iron-containing cofactor heme with high affinity in hemoproteins; CO binds the blood oxygen-carrier hemoglobin (Hb) to form carboxyhemoglobin (COHb) with an affinity 200 to 400-fold greater than oxygen, thereby diminishing oxygen delivery to tissues (11, 12). CO binding to hemoglobin stabilizes the allosteric transition from low-affinity tense state hemoglobin (Hb-T) to high-affinity relaxed state hemoglobin (Hb-R), increasing the binding affinity of CO and other diatomic ligands and further limiting oxygen binding and delivery (13). In oxygen-starved tissues, CO specifically inhibits the hemoprotein cytochrome *c* oxidase in the mitochondrial electron transport chain, uncoupling respiration and reducing energy availability (14–17). Diminished mitochondrial respiration causes damage in tissues with high energy demands, such as cardiac and neuronal tissues, contributing to long-term sequelae in survivors of CO poisoning (1, 2).

To date, no antidotal therapy for CO poisoning exists, and treatments are limited to inhalation of 100% normobaric or hyperbaric oxygen. These treatments enhance CO clearance by increasing the partial pressure of oxygen and thereby increasing the rate of CO exchange in red blood cell (RBC) hemoglobin in the lungs. With 100% normombaric oxygen, the half-life of HbCO is decreased from 320 minutes to around 74 minutes (1). Hyperbaric oxygen administration can further accelerate the CO elimination; however, there is often a several hour delay between diagnosis of CO poisoning, patient transport, and treatment, reducing the effectiveness of this strategy (7, 18, 19). The latency between diagnosis and treatment, as well as persistence of long-term sequelae in patients both with and without supplemental oxygen inhalation treatment, highlight the need for point-of-care antidotes to treat CO poisoning.

Our laboratory has specialized in a promising therapeutic strategy to treat acute CO poisoning that relies on administration of high-affinity CO scavenging agents. These scavenging agents leverage the strong interaction between CO and the Fe(II) (ferrous) heme cofactor bound to a protein or small-molecule scaffold, and tight CO binding to these agents sequesters CO away from endogenous hemoprotein sites (20–23). We first demonstrated this concept with a high CO affinity recombinant hemoprotein variant of human neuroglobin (Ngb-H64Q-CCC) to treat acute CO poisoning *in vivo* (21, 24). Infused Ngb-H64Q-CCC, which exhibits 500-fold higher CO affinity than hemoglobin, depletes the fraction of HbCO in RBCs, reverses hemodynamic collapse, restores mitochondrial function, and improves survival in an otherwise lethal murine model of acute CO poisoning. Infusion of other hemoproteins that have similar CO affinities to RBC hemoglobin, including alkylated hemoglobin and myoglobin (Mb), also improve hemodynamic outcomes and survival in murine models of acute CO poisoning, though these proteins likely function by both providing modest CO scavenging and enhancing tissue oxygen delivery (22). In addition to these hemoproteins, several small molecule-based scavengers have shown promise as CO sequestration agents in recent years (20, 23, 25, 26). Recently, Mao et al showed that treatment with the water-soluble synthetic heme mimic, hemoCD, demonstrated efficacy against CO poisoning in mice (23). Taken together, these proof-of-concept studies demonstrate the feasibility of intravenous CO scavenging to treat acute CO poisoning; however, questions remain regarding these molecules’ optimal therapeutic window.

Generally, hemoprotein-based scavengers must 1) possess a high CO binding affinity and kinetic parameters to outcompete physiological hemoprotein targets (e.g., hemoglobin, *K*_d,CO_ = 1.7 nM for Hb-R) (11); 2) exhibit selectivity for CO over oxygen, as high scavenger oxygen affinity would inhibit CO scavenging; 3) be stable to thermal and chemical degradation to prevent heme release and subsequent adverse reactivity in physiological environments (27); and 4) exhibit redox stability, *i.e.*, slow autoxidation from Fe(II)-O_2_ to Fe(III) heme, as only the Fe(II) heme readily binds CO. Additionally, as shown historically in the development of hemoglobin-based blood substitutes, intravenous addition of hemoproteins exhibit nitric oxide (NO) scavenging reactions, triggering hypertension and heme oxidation leading to release of pro-inflammatory ferric heme. Such off-target reactions lead to acute renal failure and increased mortality in pre-clinical and clinical trials (28, 29). To circumvent these off-target effects, we seek to diversify the potential pool of CO scavenging therapeutic candidates through engineering of non-globin-based hemoproteins. By fine-tuning the amino acid composition of the hemoprotein scaffold, we can optimize critical properties for CO scavenging (ligand binding parameters, redox stability, and pharmacokinetic profile) while mitigating off-target effects observed through NO reactivity.

A hemoprotein called the <u>r</u>egulator of <u>CO</u> <u>m</u>etabolism (RcoM) protein, a bacterial, non-globin transcription factor that activates aerobic CO metabolism in the presence of CO (30, 31) may serve as an ideal platform to engineer a safe, efficacious CO scavenger. Originally isolated from the soil bacterium *Paraburkholderia xenovorans*, RcoM utilizes heme to sense low environmental concentrations of CO (0.1 to 25 nM) in aerobic environments, suggesting that RcoM exhibits high CO affinity, selectivity for CO over oxygen, and heme redox stability (32). The native protein is a homodimer comprised two domains: (1) an N-terminal, sensory PAS (Per-Arnt-Sim) domain, where heme and CO binding occurs, and (2) a C-terminal, DNA-binding LytTR domain (33). Studies of the RcoM-2 paralog from *P. xenovorans* (*Px*RcoM-2) report nanomolar CO binding affinity; however, a truncate bearing only the N-terminal heme-binding domain (HBD) exhibits 16-fold higher CO binding affinity (*K*_d,CO_ = 0.25 nM for *Px*RcoM-2-HBD) (34). Importantly, oxygen binding to full-length RcoM has not been observed to date, suggesting exquisite CO selectivity. Ferrous heme in RcoM is six-coordinate; the protein binds heme via a proximal histidine (His77) and a distal methionine (Met104), the latter of which is readily replaced by CO (30, 35, 36). This Met104 residue, which saturates the heme coordination environment, may impart additional heme stability and ligand selectivity without compromising CO binding affinity.

Given the inherent characteristics of RcoM and its truncate, we hypothesize that we can devise a safe, efficacious CO scavenging hemoprotein using the RcoM platform, but several important ligand binding and stability criteria require further optimization. Herein, we present the design and characterization of a novel hemoprotein-based scaffold based on the *Px*RcoM-1 paralog as a promising CO scavenging therapeutic. We demonstrate the efficacy of this scavenger, RcoM-HBD-CCC, in a murine model of severe CO poisoning, where RcoM administration accelerates CO clearance. We further demonstrate that this scavenger elicits neither hypertension nor organ-specific toxicity upon intravenous infusion in mice due to limited reactivity with NO.

## RESULTS

### Design and recombinant expression of heme-loaded RcoM-HBD-CCC

We introduced two modifications to the native *Px*RcoM-1 protein sequence to improve CO scavenging properties. First, we removed the C-terminal, DNA-binding LytTR domain (residues 154 to 266), yielding a truncate bearing the N-terminal, heme-binding PAS domain (Figure S1a). This C-terminal domain is not involved in direct CO binding. Second, we introduced three Cys-to-Ser amino acid substitutions (C94S, C127S, and C130S) to the HBD truncate to eliminate the potential for intermolecular disulfide bond formation at high protein concentrations, as informed by previous studies (21). To facilitate isolation of high-purity recombinant protein, we appended a C-terminal 6xHis affinity tag. Initial expression of this RcoM-HBD-CCC construct, cloned into pET28a and expressed in soluBL21 *E. coli*, resulted in high protein yields (∼44-49 mg purified holoprotein/L culture) but low heme loading (16-20% based on a 1:1 ratio of protein monomer to heme cofactor, Table S1). To increase the amount of heme-loaded protein relative to total isolated protein, RcoM-HBD-CCC was co-expressed with *E. coli* ferrochelatase (*Ec*FeCH), which catalyzes the final step in heme biosynthesis. Co-expression of RcoM-HBD-CCC with *Ec*FeCH dramatically improved heme loading (102-136% heme loading, see Materials and Methods), although overall protein yields diminished (4-15 mg purified holoprotein/L culture) due to *Ec*FeCH co-expression.

### Spectroscopic properties and stability of the RcoM-HBD-CCC heme

The RcoM-HBD-CCC truncate bears a six-coordinate heme in both Fe(III) and Fe(II) oxidation states (Figure 1A and 1B, Table S2). Spectroscopic data suggest that Fe(III) heme from the native RcoM-1 homolog from *P. xenovorans* is axially coordinated by a charged thiolate (from Cys94) and neutral imidazole (from His74)(30, 35). The Fe(III) heme of RcoM-HBD-CCC is also six-coordinate, as evidenced by the positioning of the Q-band (visible) absorbance features at 537 nm and 565 nm, despite the fact that the native coordinating Cys ligand (Cys94) is replaced by a non-coordinating Ser residue. It is possible that a different protein-derived ligand occupies the sixth axial position, giving rise to the low-spin, Fe(III) spectral features. Similar observations have been reported after removing the distal ligand in some cytoglobin and neuroglobin mutants (37–40). RcoM-HBD-CCC holoprotein exhibits very high thermal stability, as measured by loss of Fe(III) heme signal, with a melting temperature value, *T*_m_, of 72 °C in phosphate-buffered saline (PBS, 10 mM, pH 7.4) (Figure 1C). Chemical reduction of the RcoM-HBD-CCC heme with sodium dithionite gives rise to an UV-Vis absorbance feature with two sharp bands at 531 nm and 562 nm, consistent with formation of a low-spin, six-coordinate heme center. Native RcoM undergoes a redox-mediated ligand switch in which the charged thiolate of Cys94 is replaced by a neutral thioether, Met104 (36). Ferrous heme coordination by Met104 is maintained in RcoM-HBD-CCC, as substitution of Met104 with a non-coordinating Leu residue (M104L) broadens the Q-bands, consistent with the formation of a five-coordinate, high-spin species (Figure S2). We note here that though the Fe(II) heme in RcoM-HBD-CCC is weakly bound by the distal Met104, for clarity we will use the term “unliganded Fe(II) heme” to describe RcoM-HBD-CCC species without an exogenous diatomic ligand bound (*i.e.*, CO, NO, and oxygen).

**Figure 1.**
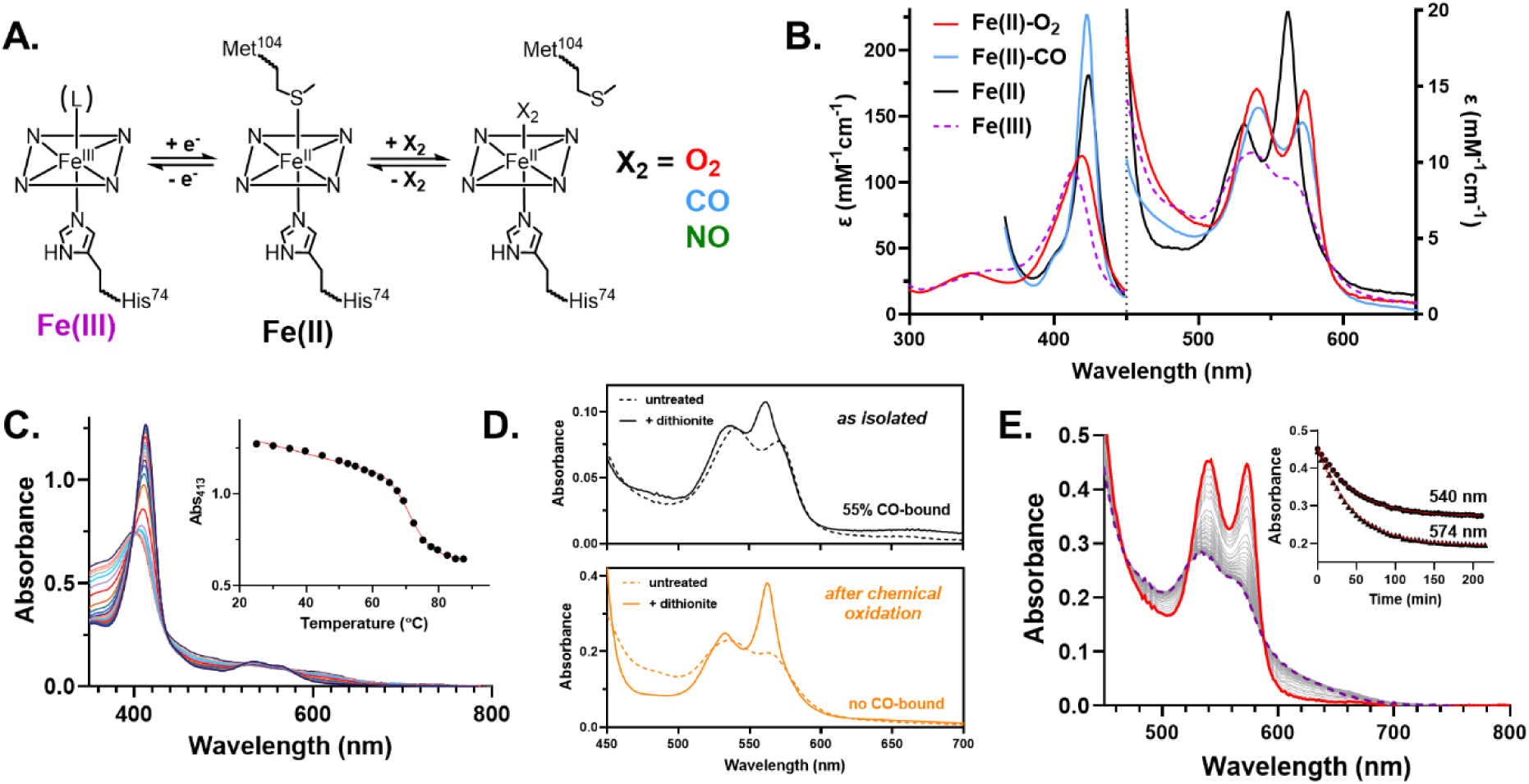
RcoM-HBD-CCC spectroscopic and stability properties All measurements carried out in phosphate buffered saline (PBS, 10 mM, pH 7.4). **(A)** Heme coordination environment for RcoM-HBD-CCC. **(B)** Electronic absorption (UV-Vis) spectra corresponding to distinct heme oxidation and ligation states for RcoM-HBD-CCC. **(C)** Thermal unfolding of RcoM-HBD-CCC bearing Fe(III) heme, as monitored by heme loss using UV-Vis spectroscopy. Inset: Decrease in absorbance at 413 nm, corresponding to dissociation of Fe(III) heme from the protein. The melting curve was fit using the Santoro-Bolen equation (red line) to determine melting temperature. **(D)** Comparison of purified RcoM-HBD-CCC UV-Vis spectra before and after chemical oxidation with excess potassium ferricyanide. Top: Complete reduction of hemoprotein and oxygen removal using sodium dithionite yields an admixture of unliganded Fe(II) and Fe(II)-CO species for homogeneous protein directly after heterologous expression and purification. Bottom: Reduction of heme using sodium dithionite after chemical oxidation yields pure unliganded Fe(II) species. **(E)** Autoxidation kinetics for Fe(II)-O_2_ RcoM-HBD-CCC heme under aerobic conditions at 37 °C. Inset: Changes in absorption at 540 nm and 574 nm, indicating decay of Fe(II)-O_2_ heme to Fe(III) heme. Curves were fit to single exponential decay functions to determine the observed rate of autoxidation, *k*_autox_ = 1.1 h^-1^ (red lines).

The ligand binding properties of RcoM-HBD-CCC differ from those of the native full-length RcoM sensor protein. We observe a majority fraction of RcoM-HBD-CCC bearing Fe(II)-CO heme (55-85% Fe(II)-CO heme; 45-15% Fe(III) heme) after isolation under typical aerobic expression conditions from *E. coli* without addition of CO to culture media (Figure 1D), an observation consistent with prior studies of RcoM truncates and full-length variants bearing the M104L substitution (30, 35, 36, 41). As with other recombinant CO scavengers, such as Ngb-H64Q-CCC (21, 39, 42), isolation of CO-bound protein during heterologous production is indicative of high CO binding affinity. Potassium ferricyanide oxidizes Fe(II)-CO RcoM-HBD-CCC, yielding Fe(III) heme (Figure 1D) and releasing heme-bound CO. This re-oxidized protein undergoes facile chemical reduction with sodium dithionite to generate the homogeneous Fe(II) (ferrous) Met104-bound species. This ferrous species also readily binds nitric oxide (NO) to yield spectroscopic features consistent with a low-spin, six-coordinate Fe(II)-NO species (Table S2). Additionally, Fe(II) RcoM-HBD-CCC forms a stable Fe(II)-O_2_ adduct not previously described for full-length or other truncated RcoM variants (Figure 1B). While the peak maxima of these Fe(II)-O_2_ features (Q-bands at 540 nm and 573 nm) are very similar to those of the Fe(II)-CO species, the band shapes and relative intensities are distinct. The RcoM-HBD-CCC Fe(II)-O_2_ adduct is relatively stable towards decomposition into Fe(III) heme and superoxide with an apparent autoxidation rate, *k*_autox_, of 1.1 h^-1^ (Figure 1E), faster than hemoglobin or myoglobin but much slower than rates reported for cytoglobin or neuroglobin (39, 43).

### CO binding parameters for RcoM-HBD-CCC

To quantify the CO affinity constant (*K*_a,CO_) of RcoM-HBD-CCC, we first sought to measure kinetic rate constants for CO binding (*k*_on,CO_) and dissociation (*k*_off,CO_). We can then compute *K*_a,CO_ as a ratio of binding association and dissociation rate constants (*K*_a,CO_ = *k*_on,CO_/*k*_off,CO_. We initially attempted to quantify *k*_on,CO_ using flash photolysis-rebinding studies; however, we observed minimal CO escape from the heme pocket after photolysis, consistent with complete geminate recombination of CO and Fe(II) heme (*vide infra*). Given that CO escape from the heme pocket is needed to quantify *k*_on,CO_, this result precluded the use of flash photolysis-rebinding. Instead, we followed CO binding kinetics for RcoM-HBD-CCC using stopped-flow electronic absorption (UV-Visible) spectroscopy, observing CO binding to RcoM-HBD-CCC on the millisecond timescale (Figure 2A). CO binding kinetics followed a single exponential function under pseudo-first order conditions (Figure 2A, *right*); changes in observed rates varied linearly as a function of CO concentration, giving rise to the second-order rate constant (*k*_on,CO_) of 4.67×10^4^ M^-1^s^-1^ (Figure 2D). In a prior stopped-flow study investigating CO binding kinetics to Fe(II) heme in a HBD truncate of the *Px*RcoM-2 ortholog, the observed binding rate constant approached a limiting value at CO concentrations of 0.5 mM in PBS, presumably due to the limitations of Met104 dissociation from heme (41). We observed no such limiting rate of CO binding; however, we note that we were unable to achieve a concentration of CO greater than 0.3 mM in solution using our stopped-flow setup.

**Figure 2.**
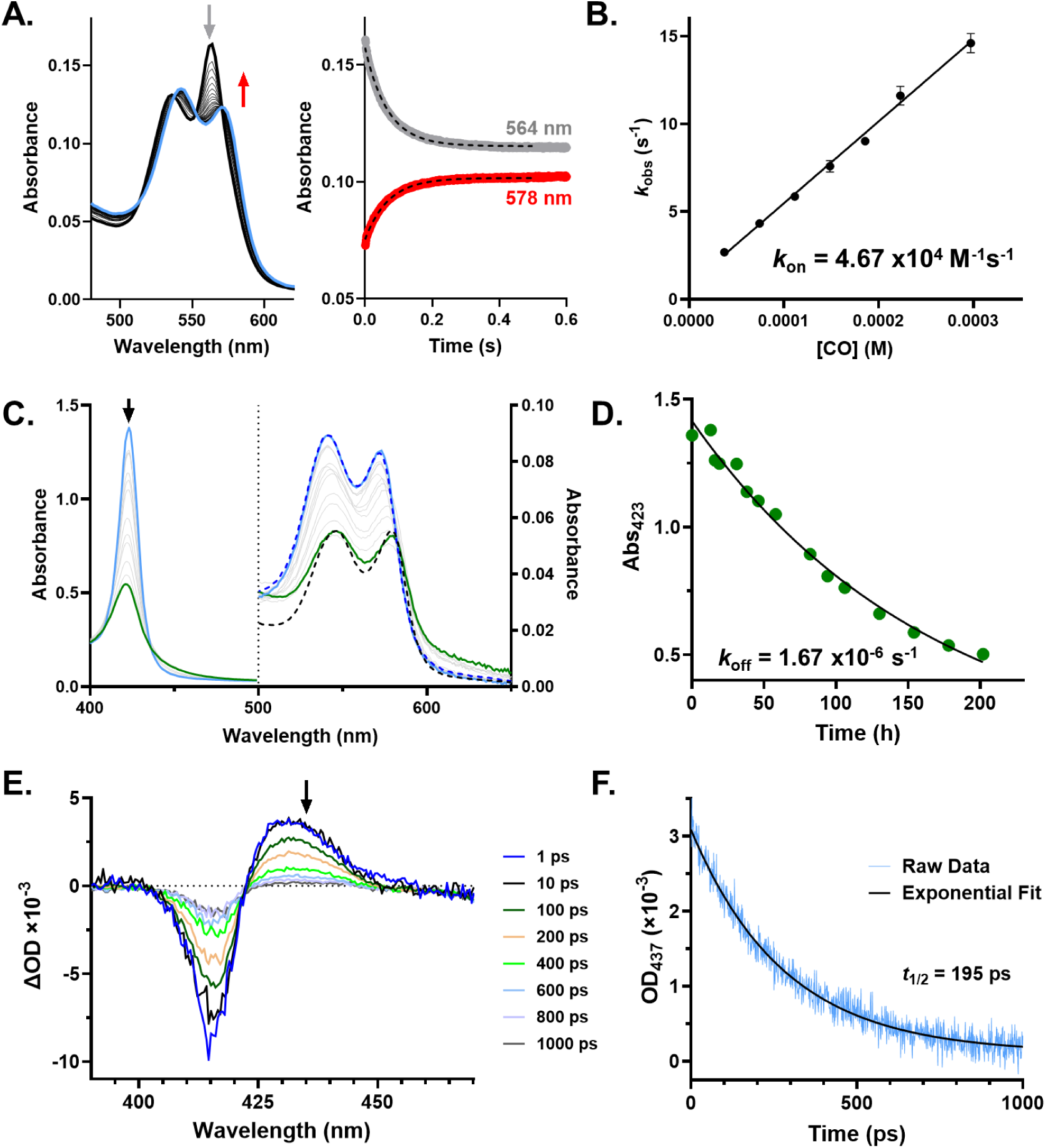
Determination of kinetic and thermodynamic parameters for CO binding to RcoM-HBD-CCC under anaerobic conditions. **(A)** (*Left*) Spectral changes in the visible (Q-band) region upon stopped-flow rapid mixing of Fe(II) RcoM-HBD-CCC (10 μM, solid black line) and CO-saturated PBS at 25 °C and 297 μM CO, resulting in formation of Fe(II)-CO RcoM-HBD-CCC (solid blue line). (*Right*) Corresponding kinetic traces following the loss of signal from Fe(II) RcoM-HBD-CCC (absorbance at 564 nm, grey line) and formation of Fe(II)-CO RcoM-HBD-CCC (absorbance at 578 nm, red line) at 25 °C. Black dashed lines depict best fits to single exponential functions for each trace. **(B)** Plot of observed pseudo first-order rate constants for CO binding (*k_obs_*) to Fe(II) RcoM-HBD-CCC as a function of solution concentration of CO at 25 °C. Each data point represents the average observed rate constant for four separate mixing events. A simple linear regression was fit yields the second-order rate constant of 4.67×10^4^ M^-1^s^-1^ for CO binding to RcoM-HBD-CCC. **(C)** Kinetics of CO dissociation from Fe(II)-CO RcoM HBD-CCC (6.6 μM) at 25 °C, as measured by replacement with NO in NO-saturated solution over the course of 8 days. Reference data for Fe(II)-CO (blue dashed line) and Fe(II)-NO (black dashed line) RcoM-HBD-CCC species are superimposed over CO displacement data in the Q-band region, normalized to highest peak intensity for each respective species. **(D)** Corresponding kinetic trace of CO dissociation, as measured by change in Soret absorbance at 423 nm as CO is replaced by NO at Fe(II) heme (green circles). The observed first-order rate constant for CO dissociation (1.67×10^-6^ s^-1^) was determined from a curve of best fit to the change in Soret absorbance intensity (black line). Transient electronic absorption spectra for Fe(II)-CO RcoM-HBD-CCC at different delay times (*t*) following CO dissociation by flash photolysis. Soret features are plotted as difference spectra where ΔOD(*t*)=Abs(*t*)-Abs(*t*_0_) and *t*_0_ is the initial Fe(II)-CO spectrum. **(F)** CO geminate rebinding kinetics as monitored by the change in Soret intensity at 437 nm following flash photolysis (blue line). The black line represents the curve of best fit to a single exponential function (*t_½_*= 195 ps; τ=281 ps).

CO dissociation from the RcoM-HBD-CCC heme is extremely slow. We measured CO dissociation kinetics under an atmosphere of anaerobic NO using UV-Vis spectroscopy by following CO replacement with NO (Figure 2C). We observed a slow isosbestic shift as pre-formed Fe(II)-CO protein is slowly converted to the Fe(II)-NO species. By fitting the change in absorption for the Fe(II)-CO Soret peak at 423 nm to a single exponential function, we obtained a first-order dissociative rate constant (*k*_off,CO_) of 1.67×10^-6^ s^-1^ (*t*_1/2_ = 115 h, Figure 2D).

Taking the ratio of experimentally determined rate constants for CO binding and dissociation, we obtain a value of *K*_a,CO_ = 2.80×10^10^ M^-1^ for RcoM-HBD-CCC, a value nearly 50-fold higher than that of Hb-R (Table 1). We also note that the observed CO dissociation rate constant for RcoM-HBD-CCC is the lowest for any characterized hemoprotein (34), and it is slow dissociation of CO from heme that primarily drives the high CO binding affinity for this engineered CO scavenger (Table 1).

**Table 1.** Summary of kinetic and thermodynamic binding parameters for oxygen and CO for select hemoproteins.

| protein | O<sub>2</sub> $k_{on}$<br>(M<sup>-1</sup>s<sup>-1</sup>) | O<sub>2</sub> $k_{off}$<br>(s<sup>-1</sup>) | O<sub>2</sub> $K_A$<br>(M<sup>-1</sup>) | CO $k_{on}$<br>(M<sup>-1</sup>s<sup>-1</sup>) | CO $k_{off}$<br>(s<sup>-1</sup>) | CO $K_A$<br>(M<sup>-1</sup>) | M<br>($K_{Co}/K_{O2}$) |
| --- | --- | --- | --- | --- | --- | --- | --- |
| RcoM-HBD-CCC | (8.2×10<sup>5</sup>)* | 5.9 | 1.4×10<sup>5</sup> | 4.7×10<sup>4</sup> | 1.7×10<sup>-6</sup> | 2.7 to 2.8×10<sup>10</sup> | 1.9×10<sup>5</sup> |
| Hb R<sup>1</sup> | 5.0×10<sup>7</sup> | 15 | 3.3×10<sup>6</sup> | 6.0×10<sup>6</sup> | 1.0×10<sup>-2</sup> | 6.0×10<sup>8</sup> | 180 |
| Hb T<sup>1</sup> | 4.5×10<sup>6</sup> | 1.9×10<sup>3</sup> | 2.4×10<sup>3</sup> | 8.3×10<sup>4</sup> | 9.0×10<sup>-2</sup> | 9.2×10<sup>5</sup> | 390 |
| StHb<sup>2</sup> | 2.4×10<sup>7</sup> | 15 | 1.6×10<sup>6</sup> | 5.1×10<sup>6</sup> | 1.3×10<sup>-2</sup> | 3.9×10<sup>8</sup> | 242 |
| Ngb-H64Q-CCC<sup>3</sup> | 7.2×10<sup>8</sup> | 18 | 3.9×10<sup>7</sup> | 1.6×10<sup>8</sup> | 4.2×10<sup>-4</sup> | 3.8×10<sup>11</sup> | 9.7×10<sup>3</sup> |
| hemoCD<sup>4</sup> | 4.7×10<sup>7</sup> | 1.3×10<sup>3</sup> | 3.6×10<sup>4</sup> | 1.3×10<sup>7</sup> | 2.5×10<sup>-4</sup> | 5.2×10<sup>10</sup> | 1.4×10<sup>6</sup> |

1.Values from Cooper, C.E. *Biochim. Biophys. Acta Bioenerg.* **1999**
2.Values from Xu, Q. et al. *J. Clin. Invest. Insight* **2022**
3.Values from Azarov, I. et al. *Sci. Transl. Med.* **2016**
4.Values from Kano, K. et al. *Inorg. Chem.* **2006**

Slow dissociation of CO from the RcoM-HBD-CCC heme is likely driven by fast and complete geminate recombination. Ultrafast transient electronic absorption spectroscopy was employed to quantify geminate CO rebinding kinetics for Fe(II)-CO RcoM-HBD-CCC. Following a 200 femtosecond pulse at 570 nm, we observe a difference absorbance spectrum characteristic of a transition from 6-coordinate, low-spin Fe(II) to 5-coordinate high-spin Fe(II) heme (Figure 2E), consistent with complete CO photolysis. Within 1 ns, the amplitude of this difference spectrum rapidly diminished, and the spectrum returned to that of Fe(II)-CO RcoM-HBD-CCC. The loss of 5-coordinate, high-spin Fe(II) signal followed a single exponential decay (*t*_1/2_=195 ps; τ=281 ps) and approached a signal plateau that was 3.5% that of the initial amplitude (Figure 2F), demonstrating rapid and nearly complete geminate recombination of CO.

### Oxygen binding parameters for RcoM-HBD-CCC

In addition to possessing a CO binding affinity (*K*_d_) in the picomolar range, RcoM-HBD-CCC exhibits remarkable selectivity for CO over oxygen. A low autoxidation rate allowed for direct quantification of the oxygen binding affinity (*K*_a,O2_) for Fe(II) RcoM-HBD-CCC using standard tonometry methods and UV-Vis spectroscopy to monitor oxygen binding (Figure S3). Using the Hill equation to fit the oxygen binding curve, we obtain an association binding constant *K*_a,O2_ = 1.4×10^5^ M^-1^ (*P*_50_ = 3.98 mmHg) for RcoM-HBD-CCC. This value is five orders of magnitude lower than that of CO, giving rise to a selectivity constant (M-value) of 1.9×10^5^ (Table 1). We used stopped-flow UV-Vis spectroscopy to quantify oxygen dissociation from Fe(II) RcoM-HBD-CCC heme in the presence of excess CO, obtaining a *k*_off,O2_ of 5.9 s^-1^ (Figure S3). With the experimental values for *K*_a,O2_ and *k*_off,O2_, we computed the oxygen association rate constant (*k*_on,O2_) for Fe(II) RcoM-HBD-CCC, obtaining a value of 8.2×10^5^ M^-1^s^-1^. Table 1 compares these values to R- and T-state hemoglobin, as well as these parameters for other published CO scavengers.

### *In vitro* CO scavenging by RcoM-HBD-CCC

RcoM-HBD-CCC effectively sequesters CO from RBC-encapsulated HbCO under aerobic and anaerobic conditions *in vitro*. To model CO scavenging in circulation, we incubated CO-saturated murine RBCs (>90% HbCO) with Fe(II)-O_2_ RcoM-HBD-CCC at equimolar final concentrations of 100 μM heme under aerobic conditions at 37 °C. Using UV-Vis spectroscopy, we followed CO transfer from encapsulated Hb to extracellular RcoM and used spectral deconvolution to quantify the fraction of CO-bound and CO-free hemoproteins in RBC and extracellular compartments (Figure 3). We observed rapid loss of 0.72 ± 0.03 equivalents of CO from RBC Hb in approximately five minutes, with a HbCO decay half-life of 27 ± 4 s. A slightly higher than expected fraction of Fe(II)-CO RcoM-HBD-CCC from CO transfer from Hb was observed (0.86 ± 0.04 equivalents), likely due to a small amount of excess CO dissolved in solution during the preparation of CO-saturated RBCs. CO transfer is slightly slowed at 25 °C under otherwise identical conditions (HbCO decay half-life of 38 ± 2 s), attributed to the decreased kinetics of ligand exchange (Figure S4A). Under these same conditions, when one equivalent of Fe(II)-O_2_ RcoM-HBD-CCC is incubated with a five-fold excess of RBC-encapsulated HbCO, nearly complete transfer of one equivalent of CO from Hb to RcoM is observed at an accelerated rate, with a HbCO decay half-life of 15 ± 2 s (Figure S4B). Under anaerobic conditions and 25 °C, we observe complete, rapid transfer of one equivalent of CO from RBC-encapsulated deoxyHb to Fe(II) RcoM-HBD-CCC (Figure S4C). Taken together, these data demonstrate that RcoM-HBD-CCC sequesters the majority of CO from HbCO within minutes in RBCs under both aerobic and anaerobic conditions.

**Figure 3.**
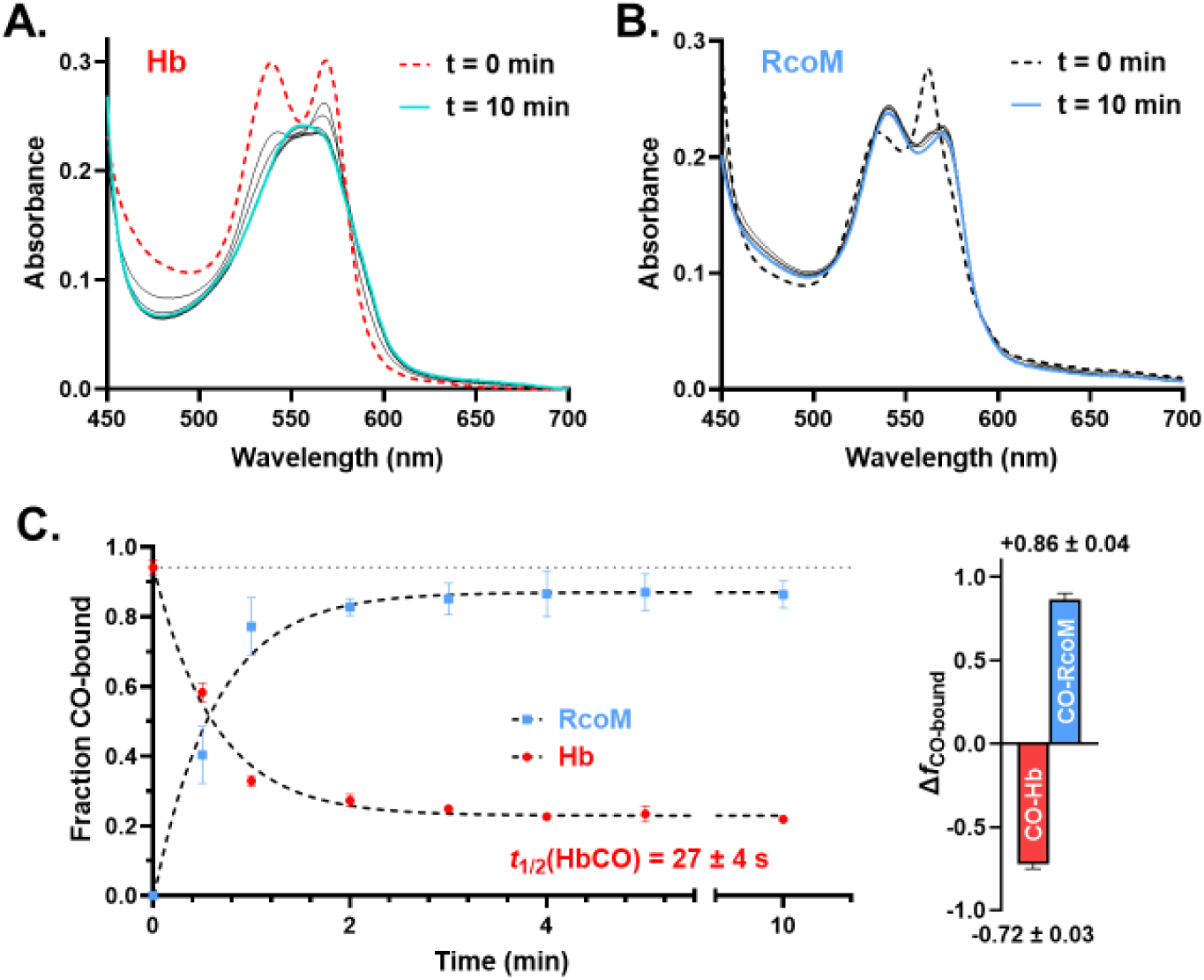
RcoM-HBD-CCC rapidly scavenges CO from RBCs *ex vivo* under aerobic conditions. CO-saturated murine RBCs (>90% HbCO, 13 μM heme) were incubated with RcoM-HBD-CCC (>99% Fe(II)-O_2_, 13 μM heme) at 37 °C for 10 min total. **(A)** Representative UV-Vis spectra for lysed RBC samples treated with RcoM-HBD-CCC *in vitro.* In the initial sample (red dashed line), Hb from RBCs is primarily (93%) CO-bound, as indicated by prominent features at 540 and 570 nm. Over the course of the 10 min experiment, these features in the isolated Hb decay over several minutes, reaching a final spectrum (solid cyan line) with peak maximum at 560 nm, a feature characteristic of unliganded Fe(II) Hb heme. **(B)** Representative UV-Vis spectra for cell-free samples containing RcoM-HBD-CCC. In the initial sample (black dashed line), prominent features characteristic of unliganded Fe(II) heme are observed at 531 nm and 562 nm. These features decay over several minutes, approaching a final spectrum (solid blue line) with peak maxima at 541 and 572 nm, characteristic of Fe(II)-CO RcoM-HBD-CCC heme. (**C**) Combined kinetic data summarizing CO transfer from RBC-encapsulated HbCO (red circles) to extracellular RcoM-HBD-CCC (blue squares). Fractions of CO-bound species were determined by spectral deconvolution. Each data point represents the average value from three technical replicates, and error bars depict ± 1 standard deviation. Dashed lines represent best fits to single-exponential functions (*k*_obs_ = 0.027 ± 0.003 s^-1^). See Materials and Methods for more details.

### *In vivo* CO scavenging by RcoM-HBD-CCC in a severe, nonlethal CO poisoning model

RcoM-HBD-CCC accelerates CO clearance from circulation in mice challenged with acute, severe CO poisoning. Using a previously established model of severe, non-lethal CO poisoning (22), we assessed hemodynamic outcomes and scavenger-dependent changes in the fraction of CO-bound hemoglobin in RBCs (*f*_HbCO_) in mice. Tracheally-intubated, anaesthetized animals under mechanical ventilation were exposed to a 3% CO gas mixture balanced with air for 1.5 min resulting in COHb levels of ∼76% in these animals. Three minutes after CO exposure, PBS vehicle or Fe(II)-O_2_ RcoM-HBD-CCC (scavenger) in PBS was administered intravenously via a jugular vein catheter at scavenger protein dosages of 255 mg/kg (15 μmol*/kg), 510 mg/kg (30 μmol/kg), or 1020 mg/kg (60 μmol/kg) and a dosing volume of 10 mL/kg body weight, or a total scavenger volume of ∼300 μL (Figure 4A). Following CO inhalation, mean arterial pressure (MAP) drops from 80-85 mmHg to ∼50 mmHg (Figure 4B, Table S3. Infusion occurs with a transient increase in blood pressure to values around 100 mmHg in both vehicle and scavenger treatment groups, followed by a stabilization to blood pressure values at or slightly above baseline within 10 min of CO exposure.

**Figure 4.**
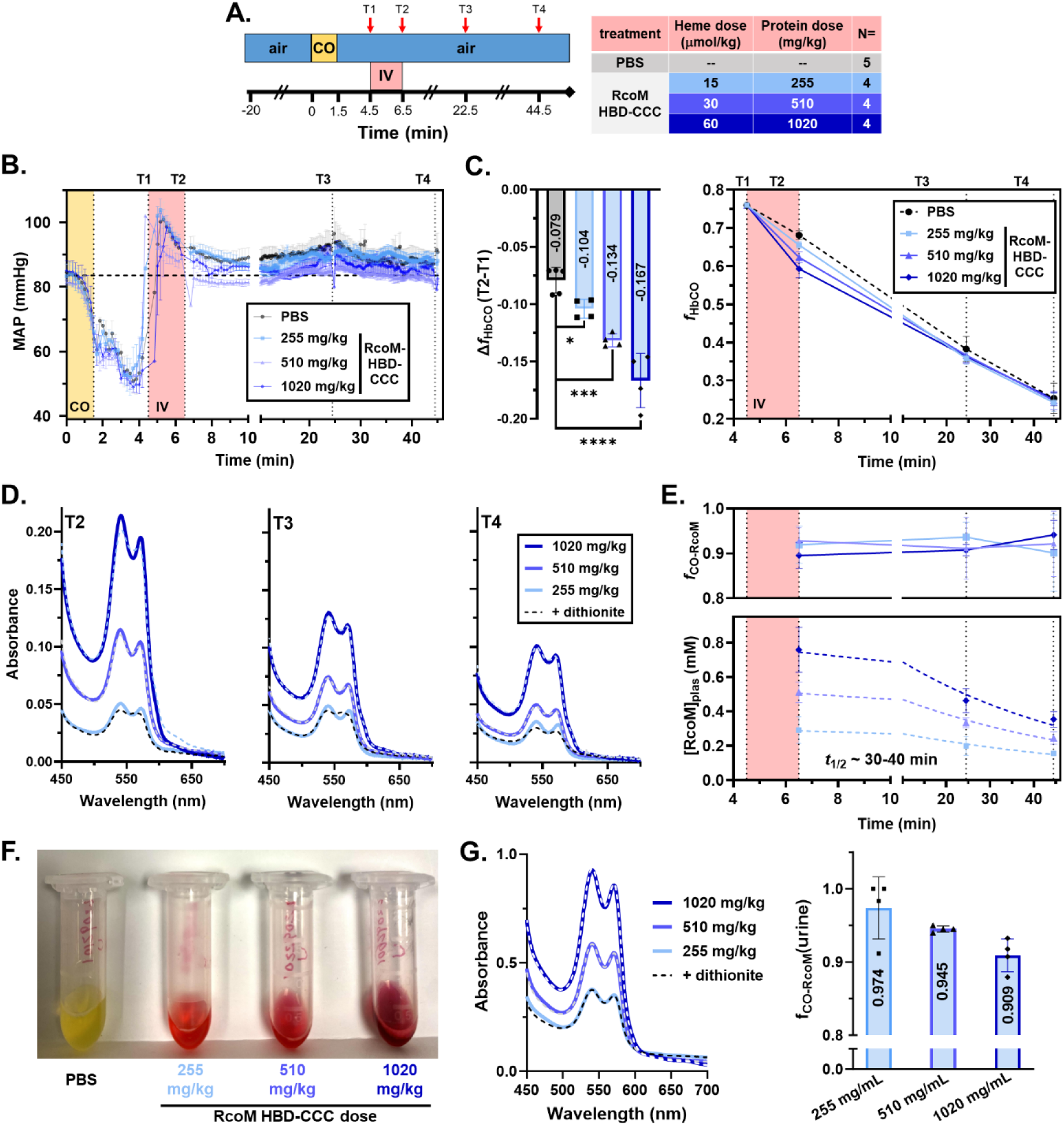
RcoM-HBD-CCC accelerates CO clearance in a murine model of severe CO poisoning. **(A)** (*left*) Experimental scheme for severe CO poisoning model in anaesthetized mice under mechanical ventilation via tracheal tube. Intravenous delivery of RcoM-HBD-CCC scavenger or PBS vehicle was carried out via catheter in the jugular vein. A carotid arterial catheter was surgically instrumented for hemodynamic monitoring and blood sampling. Red arrows denote time points for 15 uL blood draws. Body temperature was monitored and maintained at 37 °C throughout the course of the experiment. (*right*) RcoM dosage and animal numbers **(B)** Changes in mean arterial pressure (MAP) as a function of time in the severe, nonlethal CO poisoning model. The average starting blood pressure across all treatment groups is depicted by the vertical dashed line at 83.6 mmHg. **(C)** (*left*) Comparison of CO clearance, as measured by the difference in *f*_HbCO_ before and after infusion, Δ*f*_HbCO_(T2-T1), as a function of scavenger dose. Statistical significance between vehicle and treatment groups was assessed using ordinary one-way ANOVA (*, p<0.05; ***, p<0.001; ****, p<0.0001). (*right*) Changes in the fraction of circulating HbCO (*f*_HbCO_) after CO exposure and delivery of RcoM-HBD-CCC scavenger. Values for *f*_HbCO_ were quantified by spectroscopic analysis of lysed RBCs isolated from whole blood drawn at four time points throughout the experiment. To better visualize dose-dependent changes in CO clearance, all *f*_HbCO_ values were adjusted to a starting value of 0.76, the average initial *f*_HbCO_ at T1 across all treatment groups. **(D)** Representative spectroscopic data for plasma samples (diluted 40-fold in PBS) taken at different time points following RcoM-HBD-CCC infusion (T2 to T4) in the severe CO poisoning model. Spectra are compared before (solid lines) and after chemical reduction using sodium dithionite (dashed lines) to highlight the degree of CO binding. **(E)** Assessment of RcoM-HBD-CCC speciation and concentration in plasma during acute CO poisoning. Values for *f*_CO-RcoM_ (*top*) and scavenger concentration (*bottom*) were quantified by spectroscopic analysis of plasma samples isolated from whole blood drawn at four time points throughout the experiment. **(F)** Representative images of urine samples collected 45 min after acute CO exposure and subsequent infusion with scavenger. A dose-dependent increase in red color, derived from eliminated RcoM-HBD-CCC scavenger, was observed. **(G)** Spectroscopic analysis of urine samples reveal that RcoM-HBD-CCC scavenger eliminated was >90% CO-bound.

CO clearance was assessed by monitoring changes in *f*_HbCO_ in lysed RBCs from samples drawn just before scavenger infusion (T1), after infusion (T2), 22.5 min (T3), and 44.5 min (T4) after CO exposure via a carotid arterial catheter (Figure 5C). Values for *f*_HbCO_ were computed as follows:

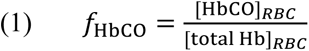

where [HbCO]_RBC_ and [total Hb]_RBC_ are quantified from spectral deconvolution of lysed RBCs. Additionally, RBCs are lysed in the presence of 20 mM sodium dithionite, which eliminates Fe(III) heme sites and scavenges heme-bound oxygen. As a result, only deoxyHb (*i.e.*, Fe(II) unliganded Hb) and HbCO species remain in the lysate, and reference data (molar absorptivity values at 1 nm increments from 450-700 nm) for these two spectroscopically distinct species are used to quantify their absolute concentrations in the admixture. The average starting value for *f*_HbCO_ in all treatment groups was 0.760 ± 0.018. In control animals infused with PBS vehicle, a decrease in *f*_HbCO_ is observed over time, consistent with both physiological clearance due to gas exchange in the lungs and loss of CO to peripheral tissues. The change in *f*_HbCO_ values before and after infusion, Δ*f*_HbCO_ (T2-T1), shows a significant, dose-dependent acceleration of CO clearance with RcoM-HBD-CCC infusion for all scavenger groups (Δ*f*_HbCO_ = −0.104 ± 0.008, −0.132 ± 0.006, −0.167 ± 0.024, respective for increasing doses) compared to the control group (Δ*f*_HbCO_ = −0.079 ± 0.011, Figure 4B). No significant differences in Δ*f*_HbCO_ were observed at subsequent time points between control and scavenger groups (Figure S5).

**Figure 5.**
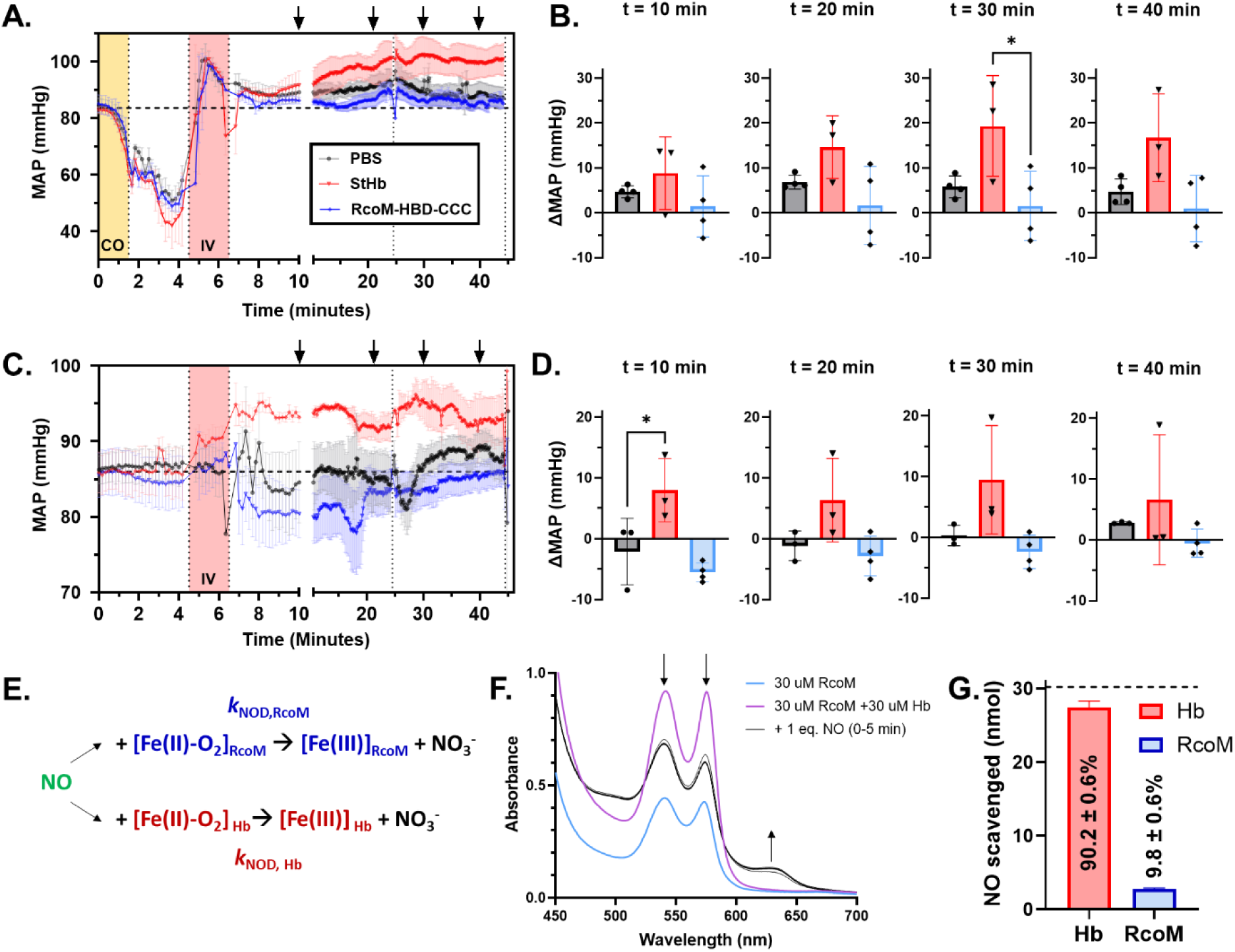
Intravenous infusion of RcoM-HBD-CCC does not elicit hypertension in mice with or without CO exposure. **(A)** Mean arterial pressure (MAP) as a function of time in the severe, nonlethal CO poisoning model with intravenous infusion of PBS vehicle (*n*=4, black), StHb at a dose of 800 mg/kg (50 μmol/kg heme; *n*=3, red), or RcoM-HBD-CCC at a dose of 1020 mg/kg protein (60 μmol/kg heme; *n*=4, blue). The average starting blood pressure across all treatment groups is depicted by the vertical dashed line at 83.6 mmHg. Black arrows denote ten-minute time intervals following CO exposure. **(B)** Highlighted values for change in MAP (ΔMAP) at specific time intervals following the initiation of CO exposure. Each data point represents the difference in blood pressure between the starting value at t=0 and ten-minute time intervals following CO exposure for each replicate. **(C)** MAP as a function of time in the severe, nonlethal CO poisoning model with intravenous infusion of PBS vehicle (*n*=3, black), StHb at a dose of 480 mg/kg protein (30 μmol/kg heme; *n*=3, red), or RcoM-HBD-CCC at a dose of 510 mg/kg protein (30 μmol/kg heme; *n*=4, blue). The average starting blood pressure across all treatment groups is depicted by the vertical dashed line at 86.0 mmHg. Note that the y-axis range in panel C differs from that in panel A. **(D)** Highlighted ΔMAP values at specific time intervals. Each data point represents the difference in blood pressure between the starting value at t=0 and ten-minute time intervals for each replicate. Statistical significance between vehicle and treatment groups was assessed using one-way ANOVA with multiple comparisons between means in each treatment group (*, p<0.05). **(E)** Reaction scheme for the estimation of the NO dioxygenation rate constant for RcoM-HBD-CCC (*k*_NOD,RcoM_) through a competition reaction with StHb. **(F)** Representative spectroscopic data for NO dioxygenation competition experiment between RcoM-HBD-CCC and StHb. 30 μM Fe(II)-O_2_ RcoM-HBD-CCC (blue line) was mixed with 30 μM Fe(II)-O_2_ StHb (magenta line) at 20 °C. A stock ProliNONOate solution was added to a final concentration of 30.2 μM NO and allowed to react for 5 min (black lines). **(G)** Final amount of NO scavenging due to NO dioxygenation by StHb (red bar, 27.4 ± 0.8 nmol) and RcoM-HBD-CCC (blue bar, 2.8 ± 0.1 nmol), as determined by spectral deconvolution (*n*=4 technical replicates).

RcoM-HBD-CCC is rapidly saturated with CO in circulation. As with RBC Hb speciation *in vitro*, we followed the speciation of intravenous RcoM-HBD-CCC scavenger in plasma, isolated from blood samples used to measure RBC HbCO described above. Spectroscopic data for diluted plasma samples are consistent with rapid CO binding by the scavenger at all doses (Figure 4D). We observed minimal changes in spectral features for plasma samples upon addition of sodium dithionite, which eliminates all oxygen, including any bound to RcoM-HBD-CCC. The lack of changes further confirm a high fraction of Fe(II)-CO RcoM-HBD-CCC vs oxygen bound, ferric or unliganded RcoM-HBD-CCC, as the former’s spectral features are unaffected by the presence of the dithionite reducing agent. Using spectral deconvolution, we determined the fraction of CO-bound RcoM-HBD-CCC (*f*_CO-RcoM_) to be greater than 90% in all scavenger treatment groups at all time points (Figure 4E, *top*). Total scavenger concentrations in plasma directly correlate with dosing; elimination half-lives for each group occur between 30-40 min, though we note that we did not follow to complete elimination from plasma here (Figure 4E, *bottom*).

RcoM-HBD-CCC is rapidly excreted in the urine. As with plasma scavenger concentrations, the concentration of RcoM-HBD-CCC eliminated in the urine increases linearly with the dose administered in this model (Figure 4F and 4G). A urine sample was isolated from each animal ∼45 min following exposure to CO. Spectral features for RcoM-HBD-CCC recovered from urine were consistent with those observed in plasma samples. Moreover, spectral deconvolution of urine samples after treatment with sodium dithionite reveal greater than 90% CO-bound RcoM-HBD-CCC in all treatment groups, indicating RcoM-HBD-CCC maintains CO ligation through facile renal clearance and excretion in urine.

### RcoM-HBD-CCC NO scavenging kinetics mitigate vasoconstriction *in vivo*

Unlike hemoglobin and modified hemoglobins (22), infusion of Fe(II)-O_2_ RcoM-HBD-CCC does not appear to elicit an increase in blood pressure in the context of severe, nonlethal CO poisoning as indicated in Figure 4B. The hemodynamic parameters for animals treated with RcoM-HBD-CCC (1020 mg/kg) compared to animals treated with a similar dose of cell-free, 2,3-diphosphoglycerate stripped human hemoglobin (StHb, 800 mg/kg, 50 μmol/kg heme) under analogous CO poison conditions above are consistent with this observation (Figure 5A,B). For all animals in all treatment groups, a transient decrease in MAP to a minimum value of 40-50 mmHg was observed 3-4 min after the start of CO exposure (Figure 5A, Table S3), consistent with what was observed in Figure 4B. Likewise, an increase in MAP was observed just prior to infusion, likely to due to a combination of adrenergic response and introduction of fluid (44–46). During the infusion period, animals in all groups (including PBS vehicle) exhibit a rapid, transient increase in MAP to a maximum value of ∼100 mmHg, followed by a decrease to pressure values just above basal values over a ∼4 min period following infusion. In animals administered RcoM-HBD-CCC or PBS vehicle, MAPs stabilized just above basal values for the duration of the experiment. In contrast, MAPs increase by ∼15 mmHg in animals administered StHb between *t*=8 min and *t*=20 min, and this hypertensive effect persists for the duration of the experiment.

To eliminate CO exposure as a potential contributing factor, we administered StHb or RcoM-HBD-CCC to healthy mice; only StHb elicited a hypertensive effect after infusion (Figure 5C,D). As with the severe, nonlethal CO poisoning model, these animals were mechanically ventilated, equipped with a venous drug delivery port, and MAP continuously monitored. However, these animals were not exposed to CO and instead inhaled medical air throughout the experiment. For this cohort of animals, all infusion treatment groups exhibited stable blood pressure values of 84-86 mmHg prior to infusion. After infusion of RcoM-HBD-CCC (510 mg/kg protein, 60 μmol/kg heme) or PBS vehicle, animals exhibited a small, transient decrease in blood pressure compared to baseline (ΔMAP= −5.5 ± 1.3 mmHg for RcoM-HBD-CCC and −2.1 ± 4.5 mmHg for PBS at *t*=10 min), followed by a return to near-basal MAP values by *t*=40 min. In contrast, healthy animals administered StHb (480 mg/kg) exhibited an immediate, significant increase in blood pressure post infusion that was sustained through the course of the experiment (*t=*40 min). This divergence in the MAP response between treatment groups in healthy animals directly parallels the effect observed in CO-poisoned animals. Taken together, these data suggest that Fe(II)-O_2_ RcoM-HBD-CCC does not elicit hypertension upon intravenous infusion, an unexpected result as hypertension is well established in animals administered StHb due to rapid NO dioxygenation and consumption (47–49).

To account for the MAP difference in healthy animals, we have determined that the rate constant for NO dioxygenation (*k*_NOD_) for Fe(II)-O_2_ RcoM-HBD-CCC is ten-fold lower than that of StHb. NO dioxygenation kinetics for oxyferrous hemoproteins are exceedingly rapid (*k*_NOD,Hb_ = 6 to 8 ×10^7^ M^-1^s^-1^) and difficult to measure, even with a stopped-flow instrument (50–52). To circumvent this limitation, we carried out a competition experiment in which equimolar amounts of Fe(II)-O_2_ RcoM-HBD-CCC, Fe(II)-O_2_ StHb, and NO were mixed in a sealed cuvette (Figure 5E). As one equivalent of NO reacts with one of Fe(II)-O_2_ heme to make Fe(III) heme, the final fraction of Fe(III) heme for each protein directly correlates to the amount of NO scavenged. Spectroscopic changes were rapid and reached final equilibrium well before the final spectrum at 5 minutes (Figure 5F). Spectral deconvolution of the final reaction mixture was facilitated by spectroscopic differences between the two Fe(III) species (Figure S6). The final distribution of Fe(III) species was 90.2 ± 0.6% Hb and 9.8 ± 0.6% RcoM-HBD-CCC (Figure 5G, Table S5). This final distribution also reflects the relative rates between RcoM-HBD-CCC (*k*_NOD,RcoM_) and StHb (*k*_NOD,Hb_), given that NO dioxygenation is irreversible and fast for both proteins. We therefore estimate a value of *k*_NOD,RcoM_ of 6-8 ×10^6^ M^-1^s^-1^ from known values of *k*_NOD,Hb_.

### RcoM-HBD-CCC safety and toxicity studies in healthy mice

Healthy (non-CO poisoned) mice tolerated intravenous infusion of RcoM-HBD-CCC without noticeable organ injury. Fe(II)-O_2_ RcoM-HBD-CCC (170 mg/kg) was administered via tail vein catheter to a separate cohort of healthy, non-CO poisoned mice. All mice administered RcoM-HBD-CCC (*N*=3) and PBS vehicle (*N*=6) survived to the end of the 48-hour monitoring period, and no significant differences in animal behavior were observed. No accumulation of RcoM-HBD-CCC in tissue (lung, liver, spleen, or kidney) was observed after the 48-hour observation period by Western blot against anti-6xHis antibodies, nor were plasma biomarkers for liver or kidney function elevated compared to animals administered PBS vehicle, demonstrating minimal organ-specific toxicity (Figure S7).

## DISCUSSION

In this study, we engineered a variant of the microbial CO-sensing transcription factor, RcoM, with ideal properties for a pharmaceutical CO antidote. We hypothesized that RcoM, which utilizes heme to sense low environmental concentrations of CO in aerobic environments, would act as an ideal platform to design a selective, high-affinity CO scavenger. Consistent with this hypothesis, we identified a truncated variant of RcoM, RcoM-HBD-CCC, that exhibits extremely high affinity for CO (*K*_d,CO_ = 37 pM), but only modest affinity for O_2_ (*K*_d,O2_ = 7.1 μM). In contrast to previously reported hemoprotein-based CO scavengers, which are globin-based (21, 22), the 17 kDa RcoM-HBD-CCC likely adopts a tertiary structure consistent with a PAS domain. The ∼100 amino acid PAS domain, found in all kingdoms of life, contains a highly conserved structural motif comprised of a five-stranded β-sheet with intervening α-helices that together form a ligand binding pocket that can be specifically modified to accommodate a wide variety of ligands (53). In the case of RcoM, this binding pocket bears the iron-containing cofactor, heme (30, 31). As in native RcoM, heme stably and irreversibly binds to RcoM-HBD-CCC under physiological conditions, as evidenced by high thermal stability (T_m_ = 72 °C). Unlike the globin-based CO scavengers, the heme iron in RcoM-HBD-CCC remains coordinatively saturated in both ferric and ferrous oxidation states (Figure 1A), which likely contributes to protein stability, relatively slow autoxidation, diatomic ligand selectivity, and lack of organ-specific toxicity.

The exceedingly slow dissociation of CO from Fe(II) heme drives high CO binding affinity and selectivity for RcoM-HBD-CCC. Our experimental data suggest a first-order rate constant of 1.67×10^-6^ s^-1^ for CO dissociation from Fe(II) RcoM-HBD-CCC. This value for RcoM-HBD-CCC is consistent with a dissociation rate constant previously measured for the heme-binding domain for the *Px*RcoM-2 ortholog (*k*_CO,off_ = 3.5×10^-6^ s^-1^) (34). As with *Px*RcoM-2, we find that slow dissociation for RcoM-HBD-CCC is likely attributed to fast (ps) and almost complete geminate recombination (Figure 2E). For a majority of hemoproteins, geminate recombination is readily observed for O_2_ and NO, but to a far lesser extent for CO (54). In fact, fast geminate recombination with CO has only been characterized in a handful of hemoproteins, including the other microbial CO-sensing transcription factor, CooA (55–57), the PAS-containing microbial oxygen sensor, DosP (58), truncated bacterial hemoglobin (59), and Met80 variants of the electron transfer protein cytochrome *c* (60). Extensive studies of these proteins using ultrafast ligand rebinding spectroscopy, high-resolution structural data, mutagenesis, and molecular dynamics simulations reveal that the distal heme pocket positionally restricts CO in an orientation perpendicular to the heme plane (54). These positional restrictions, which may be ascribed to steric or distal pocket rigidity, lower the barrier for CO re-binding such that re-binding is favored over diffusion out of the distal heme pocket. In proteins that do not exhibit significant CO geminate recombination, such as myoglobin, CO readily migrates to a position parallel to the heme plane upon photodissociation from ferrous heme, thereby favoring CO release instead of re-binding (61). While no high-resolution structural data exist for RcoM to date, we can infer from the above studies that RcoM must represent an extreme case in terms of hydrophobicity, solvent access, and/or positional restriction that favors barrierless CO recombination with heme and near-irreversible CO binding (34, 41). We speculate that such a heme pocket likely evolved to maximize sensitivity towards CO while preventing off-target activation by oxygen in aerobic niches of CO-oxidizing bacteria. From a therapeutic standpoint, this characteristic improves the CO scavenging activity, while also limiting heme-iron oxidation, protein oxidative or thermal denaturation, and NO scavenging, especially in the presence of CO.

The kinetic ligand binding parameters for RcoM-HBD-CCC enable oxygen delivery and fast, irreversible CO scavenging in the context of acute CO poisoning. Comparing kinetic binding parameters for oxygen and CO, we find that oxygen binding and dissociation at the RcoM-HBD-CCC heme are faster than CO binding and dissociation. Given that oxygen escapes the heme pocket five times faster than CO, we reasoned that RcoM-HBD-CCC could be administered bearing Fe(II)-O_2_ heme in the context of CO poisoning. At or below oxygen tensions of ∼20 mmHg, there will be rapid net release of oxygen from RcoM-HBD-CCC (*t*_1/2_ for oxygen dissociation is 0.1 s), allowing for stoichiometric delivery of one equivalent of oxygen for every RcoM-HBD-CCC heme. After oxygen release, unliganded RcoM Fe(II) heme sites are free to irreversibly bind and sequester CO in circulation or provide further oxygen transport and delivery depending on the relative concentrations of CO and oxygen in plasma. Consistent with these kinetic parameters, mixing cell-free Fe(II)-O_2_ RcoM-HBD-CCC with RBC-encapsulated HbCO in a 1:1 ratio *in vitro* resulted in ∼75% transfer of CO from Hb to RcoM within 5 minutes of mixing (Figure 3). This transfer improves to 100% under strictly anaerobic conditions (where unliganded Fe(II) RcoM-HBD-CCC is used) and suggests that RcoM-HBD-CCC should act as an effective CO scavenging agent *in vivo* during acute CO poisoning.

RcoM-HBD-CCC accelerated CO clearance in a severe, murine model of acute, inhaled CO poisoning. We compared the fraction of RBC HbCO before and after infusion of scavenger (or PBS vehicle), Δ*f*_HbCO_, over the course of a 45 min period following CO exposure (Figure 4). Immediately following infusion, we observed a dose-dependent increase in the magnitude of Δ*f*_HbCO_. Even at the smallest dose of RcoM-HBD-CCC administered (255 mg/kg), we observe a quantifiable increase in the magnitude of Δ*f*_HbCO_ compared to PBS vehicle. No significant differences in Δ*f*_HbCO_ were observed at subsequent time points between control and scavenger groups; however, prior data suggest that CO clearance just following acute CO exposure is critical for survival in lethal animal models of acute CO poisoning (21, 22). Consistent with *in vitro* kinetics, which suggest rapid, irreversible CO binding by RcoM-HBD-CCC, the scavenger was >90% CO-bound in plasma immediately following infusion and throughout the course of the experiment (Figure 4). Roughly one hour after infusion, RcoM-HBD-CCC was identified at high concentration in Fe(II)-CO bound form in the urine. These results are consistent with rapid binding, sequestration, and elimination of CO by the RcoM-HBD-CCC protein.

Remarkably, intravenous infusion of RcoM-HBD-CCC bearing Fe(II)-O_2_ heme did not lead to an increase in MAP in healthy animals nor in animals exposed to CO. This lack of vasoreactivity gives RcoM-HBD-CCC a unique advantage over other globin-based hemoproteins, which induce significant hypertension upon intravenous infusion (47–49). In cell-free oxyferrous globins, this vasoconstriction is primarily driven by depletion of tonic NO levels through fast and irreversible NO dioxygenation (51, 52, 62, 63). Several studies from Olson and colleagues reveal a strong correlation between the NO dioxygenation rate (*k*_NOD_) and the extent of hypertension induced upon intravenous infusion. For example, recombinant human hemoglobin (rHb0.1) with *k*_NOD_ = 5.8 × 10^7^ M^-1^s^-1^ gave rise to an increase in MAP of 28.4 ± 5.7 mmHg when infused in rats at a dose of 350 mg/kg, while a Hb variant (rHb4) with 30-fold lower NO dioxygenation rate (*k*_NOD_ = 2 × 10^6^ M^-1^s^-1^) only gave rise to MAP increase of 7.6 ± 1.0 mmHg, roughly a four-fold lower effect compared to that of rHb0.1 (62). We determined the rate of NO dioxygenation for RcoM-HBD-CCC to be ∼10% that of StHb (*k*_NOD,RcoM_ = 6-8 × 10^6^ M^-1^s^-1^). Given the strong correlation between *k*_NOD_ and vasoconstriction, we conclude that the lack of vasoactivity observed following infusion of RcoM-HBD-CCC primarily derives from diminished NO scavenging through the dioxygenation reaction. The discovery of an intravenously infused hemoprotein that does not trigger hypertension represents a significant advancement in the field of hemoprotein therapeutics, and we are directing future work towards the development of RcoM variants as potential non-globin-based blood substitutes.

RcoM-HBD-CCC administered at a screening dose was well-tolerated by healthy (non-CO poisoned) animals. Over the course of a 48-hour observation period, healthy mice administered Fe(II)-O_2_ RcoM-HBD-CCC (170 mg/kg) exhibited normal behavior. Organ-specific blood chemistry markers for liver and kidney damage were nominal after 48 hours, and these results were comparable to those observed in preliminary safety studies of StHb (22). Western blotting of organ tissues using a 6xHis specific antibody indicates minimal accumulation of scavenger in lung, liver, spleen, and kidney after 48 hours. These results represent a significant improvement in safety and tolerability over Ngb-H64Q-CCC, which was well-tolerated in CO-poisoned animals, but whose safety was not investigated in healthy, non-CO poisoned animals (21). Further, administration of Ngb-H64Q-CCC bearing Fe(III) heme caused kidney damage, suggesting a mechanism involving adverse heme reactivity, such as heme release or heme-facilitated reactive oxygen species generation (64–66). For RcoM-HBD-CCC, the distal heme pocket seems to preclude such oxidative reactivity, and the protein was safely excreted into the urine within hours of intravenous infusion in the absence of CO. Further dose range finding studies to assess the complete therapeutic window for RcoM-HBD-CCC are currently in progress.

In conclusion, we have successfully engineered a high-affinity, CO scavenging hemoprotein, RcoM-HBD-CCC, based on the heme-binding domain of the bacterial CO sensor, RcoM. Ligand binding kinetics and reactivity data reveal remarkable selectivity for CO over oxygen, as well as limited reactivity towards NO and hydrogen peroxide. These biochemical properties are consistent with a distal heme pocket that evolved to facilitate exclusive interactions between CO and heme through hydrophobicity, restricted solvent access, and/or positional restriction of heme-bound ligands. Intravenous infusion of RcoM-HBD-CCC during acute inhaled CO poisoning accelerated clearance of CO from circulating RBC Hb, and CO-bound scavenger was rapidly excreted in urine. The biochemical selectivity of RcoM-HBD-CCC translated to minimal off-target reactivity in heathy animals and in the context of CO poisoning. Remarkably, RcoM-HBD-CCC infusion did not induce hypertension, nor did RcoM induce organ-specific toxicity. Taken together, these data demonstrate that RcoM-HBD-CCC may act as a safe and efficacious treatment for acute CO poisoning.

## MATERIALS AND METHODS

### Materials

All chemicals were acquired from commercial suppliers unless otherwise stated. Gases, including argon, nitrogen, carbon monoxide (chemical pure grade, 99.5%), nitric oxide (chemically pure grade, 99%; bubbled through 1 M NaOH prior to use), and 30,000 ppm carbon monoxide (3% CO, 21% oxygen, 76% nitrogen) were obtained from Matheson Tri-Gas, Inc. All experiments were performed in 12 mM phosphate-buffered saline (PBS), pH 7.4, unless otherwise stated. All pH measurements were performed with a Fisherbrand Accumet pH meter equipped with a Hamilton MiniTrode electrode.

### Design and cloning of expression constructs

DNA encoding the *Px*RcoM-1 heme binding domain (residues 1-154) with C94S mutation and a C-terminal 6xHis tag (RcoM-HBD-C94S) was cloned into pET28a under control of the T7 promoter site using Gibson assembly cloning. The coding sequence was codon optimized for expression in *E. coli*. Primers (Integrated DNA Technology, Table S6) and reaction conditions for PCR amplification were designed using the NEBuilder tool, and linearization by PCR amplification was carried out according to the manufacturer’s instructions using the Q5 High-Fidelity DNA Polymerase (New England Biolabs). Linearized gene fragment for the RcoM HBD-C94S insert and background pET28a plasmid were incubated with Gibson Assembly Master Mix (New England Biolabs) for 15 min at 50 °C before transformation into chemically competent NEB 5-alpha *E. coli*. C127S and C130S substitution mutations were subsequently introduced to sequence-verified plasmid isolated from a single-colony transformant from Gibson assembly (Monarch Plasmid miniprep kit, New England Biolabs) using Q5 Site-Directed Mutagenesis Kit (New England Biolabs) and primers listed in Table S6. PCR primers and mutagenesis reactions were designed and carried out according to the NEBaseChanger online tool. Synthetic DNA synthesis and cloning for the bicistronic RcoM-HBD-CCC + *Ec*FeCH expression vector was generated by GenScript Inc. DNA sequences for RcoM-HBD-CCC bearing a C-terminal 6xHis tag and the coding sequence for *Ec*FeCH were codon optimized for expression in *E. coli*. The DNA sequence for RcoM-HBD-CCC-M104L was generated via site-directed mutagenesis using the RcoM-HBD-CCC + *Ec*FeCH plasmid background. PCR primers (Table S6) and mutagenesis reactions were designed and carried out according to the NEBaseChanger online tool.

### Heterologous expression and purification

Initial expression of RcoM-HBD-CCC alone (pET28a-based) and co-expression of RcoM-HBD-CCC and *Ec*FeCH (pETDuet1-based) were carried out in *E. coli* strain SoluBL21 (Genlantis/AMS Bio). Homogeneous protein isolated from these expression strains was utilized for biochemical and in vitro experimentation. For animal experiments, the pETDuet1-based RcoM-HBD-CCC + *Ec*FeCH plasmid was transformed into *E. coli* strain ClearColi, a BL21(DE3)-derived *E. coli* strain that produces altered lipopolysaccharide that does not elicit TLR4/MD2-mediated immune response in cultured human cells (67). For both strains, the following growth, induction, and isolation protocol was utilized except for the identity of antibiotic, which differed for pET28a-based vectors (50 μg/mL kanamycin) and pETDuet1-based vectors (50 μg/mL ampicillin or carbenicillin).

Dense overnight culture (grown in 50 mL Luria broth + antibiotic at 37 °C with shaking at 220 rpm for 16 h) was added to 1 L terrific broth liquid media with antibiotic and incubated at 37 °C with shaking at 220 rpm until OD_600_ reached 0.8-1 (≈ 2 h). Protein expression was induced by addition of isopropyl 1-thio-β-D-galactopyranoside to a final concentration of 1 mM. At the time of induction, flasks were also charged with δ-aminolevulinic acid to a final concentration of 0.4 mM to facilitate heme biosynthesis. Cultures were incubated at 30 °C with shaking at 220 rpm for 20-22 h. After growth, cells were harvested by centrifugation at 4,500 *g* for 10 min at 4 °C and stored at −20 °C until lysis and purification.

All manipulations of protein were carried out on ice or at 4 °C. Cells were resuspended in low imidazole buffer (50 mM phosphate, pH 8.0, 50 mM NaCl, 10 mM imidazole) and lysed via sonication with stirring for a total of 30 min (4 s on, 4 s off). Crude lysate was centrifuged at 17,000 *g* for 45 min to remove insoluble lysate components. Soluble lysate was then immediately loaded onto pre-packed HisTrap FF columns (Cytiva Life Sciences) and washed with low imidazole buffer until absorbance at 280 nm reached a stable baseline. Non-specifically bound proteins were removed with a wash at 10% high imidazole buffer (50 mM phosphate, pH 8.0, 50 mM NaCl, 300 mM imidazole) until absorbance at 280 nm reached a stable baseline. RcoM-HBD-CCC was eluted in 100% high imidazole buffer, red fractions pooled, and homogeneous protein buffer exchanged to 10 mM phosphate-buffered saline (PBS) with at least three successive rounds of concentration and dilution using a 10 kDa molecular weight cutoff spin concentrator at 4 °C (Millipore-Sigma). Final protein aliquots in PBS were stored at −80 °C. Protein purity for each batch was assessed to be >95% homogeneous based on quantitation of band stained with Coomassie blue after separation by SDS-PAGE (Figure S1C).

### Quantitation of heme cofactor loading

Total protein content in homogeneous (>95% pure) protein samples were determined using the Pierce 660 nm assay (Pierce-Thermo Fisher) using bovine serum albumin as a calibration standard. Heme content was assessed using the pyridine hemochromagen assay (68). Heme loading in individual protein samples was calculated as the concentration of heme divided by the concentration of total protein.

### Manipulation of heme oxidation and ligation

Spectroscopic investigation of each protein revealed that a significant fraction of protein isolated after heterologous expression bears Fe(II)-CO heme. To generate homogeneous, CO-free protein, RcoM-HBD-CCC samples were reacted with potassium ferricyanide (Sigma Aldrich, 99%) to convert Fe(II)-CO heme to Fe(III) heme. Solid potassium ferricyanide was directly dissolved in at least 50-fold molar excess relative to heme and allowed to react for 2 h at room temperature. After oxidation, protein samples were eluted through a gravity size exclusion/desalting column (Econo-Pac 10DG, Bio-Rad) pre-equilibrated with PBS. Chemical reduction of homogeneous Fe(III) hemoprotein was accomplished by dissolving solid sodium dithionite (Acros Organics, 85%) into protein solution in at least 10-fold molar excess relative to RcoM-HBD-CCC heme. CO binding was accomplished by direct injection of CO gas into the headspace of a septum-sealed glass cuvette. Fe(II)-O_2_ RcoM-HBD-CCC was generated by passing dithionite-reduced protein through a gravity size exclusion column (Econo-Pac 10DG, Bio-Rad) pre-equilibrated with PBS on the benchtop.

### Electronic absorption (UV-Vis) spectroscopy and kinetics

Electronic absorption (UV-Vis) spectroscopy was carried out using a Cary-50 or Cary 100 spectrophotometer (Agilent) equipped with a Peltier temperature controller. Stopped-flow kinetics data were recorded using a thermostatted SX-20 stopped-flow instrument fit with a direct-mount photodiode array and using the Pro-Data SX software (version 2.5.1852.0, Applied Photophysics, Ltd.). Unless otherwise stated, steady-state spectroscopic measurements were carried out in 1 cm optical path length quartz cuvettes with 1.4 mL internal volume, black walls, and septum-sealed screw top (Starna Scientific, Ltd.).

### Thermal unfolding

Thermal denaturation of was monitored by loss of RcoM-HBD-CCC Fe(III) heme signal at 413 nm. Samples (10 μM in heme) in PBS were allowed to equilibrate at each temperature for 5 min before recording spectral data.

### Determination of binding association rate constant *k*_on,CO_

The second-order rate constant for CO binding to Fe(II) RcoM-HBD-CCC heme was determined by anaerobically mixing Fe(II) RcoM-HBD-CCC with varying concentrations of dissolved CO(g) under pseudo-first order conditions and monitoring observed rates using stopped-flow UV-Vis spectroscopy. All solutions were made in septum-sealed glass serum vials, and liquid transfers were carried out using gas-tight syringes. Approximately 500 μL of Fe(III) RcoM-HBD-CCC in PBS buffer was placed in a sealed serum vial and the headspace purged with Ar_(*g*)_ for 20 min before addition of 10 mM sodium dithionite in anaerobic PBS to a final concentration of 20 μM Fe(II) RcoM-HBD-CCC. This solution was directly loaded into one of two sample syringes in the stopped-flow apparatus.

A 20 mL solution of 10 mM sodium dithionite in anaerobic PBS was bubbled for 1 h with CO_(*g*)_ to generate an anaerobic stock CO-saturated solution. Solution concentration of CO was determined by mixing stock CO-saturated PBS with deoxygenated hemoglobin at sub-stoichiometric levels of CO relative to hemoglobin heme. Spectral deconvolution was employed to assess the admixture of CO-free and CO-bound hemoglobin and thereby quantify the solution concentration of CO. This method was employed three times throughout the stopped-flow experiment, and an average of these concentrations was used as the reported concentration. The solution concentration of CO was varied by diluting CO-saturated stock with anaerobic PBS containing 10 mM sodium dithionite in a gas-tight syringe. Diluted CO solutions were then immediately loaded into the second sample syringe in the stopped-flow apparatus.

RcoM-HBD-CCC and CO solutions were then mixed in a 1:1 ratio in the stopped-flow mixing chamber and spectral features recorded for up to 10 s post-mixing. Changes in spectral features at 564 nm and 578 nm were fit to single-exponential decay functions to determine observed pseudo-first order rate constants, and final rate constant values were reported as the average of fitted rate constants from at least four reactions at each CO concentration.

### Determination of binding dissociation rate constant *k*_off,CO_

Homogeneous RcoM-HBD-CCC bearing Fe(II)-CO heme was generated by reacting chemically oxidized, Fe(III) RcoM-HBD-CCC (∼1 mM in heme) with excess solid sodium dithionite in a 1 mL screw-top glass vial with a septum cap. After reaction for 5 min at room temperature, the vial headspace was purged with CO_(*g*)_ and allowed to react for an additional 5 min. Complete CO binding to RcoM-HBD-CCC heme was assessed by UV-Vis spectroscopy, and excess CO was removed by passing the protein through a gravity size exclusion/desalting column (Econo-Pac 10DG, Bio-Rad) pre-equilibrated with 100 mM phosphate buffer, pH 7.4. Homogeneous Fe(II)-CO RcoM-HBD-CCC was diluted to a final heme concentration of 6 μM in a septum-sealed cuvette and headspace purged with Ar_(*g*)_. After allowing the cuvette to equilibrate to 25 °C in the spectrophotometer cell holder, a 20 mM stock solution of ProliNONOate (Cayman Chemical) in anaerobic 10 mM NaOH was added to the sealed cuvette using a gas-tight syringe to a final concentration of 1 mM NONOate (∼1.8 mM NO in solution). NO_(*g*)_ was periodically added to the headspace of the sealed cuvette to mitigate any NO leakage over time. Changes in spectral features consistent with replacement of CO with NO at Fe(II) heme were monitored for a total of 230 h, and the first-order rate constant for CO dissociation (*k*_off,CO_) was estimated from the first-order exponential decay function fit from the change in absorbance at 423nm.

### Ultrafast transient absorption spectroscopy

All samples were prepared to eliminate oxygen and to be equilibrated with one atmosphere of CO (so that CO in solution is about 1 mM)(69). Protein heme reduction was carried out under anaerobic conditions by initially purging sample vials and cuvettes with Ar_(*g*)_. RcoM-HBD-CCC samples were reduced with sodium dithionite (Sigma, 25 mM final concentration) and then diluted with CO-saturated PBS in a septum-capped, 1 mm pathlength cell cuvette so that the optical density was ∼1 at 420 nm. The cuvette headspace was further degassed with CO_(*g*)_ before measurements to maintain CO levels at 1 atmosphere.

The experimental system is based on a femtosecond PHAROS Yb:KGW amplifier generating 200 fs pulses at 1030 nm (Light Conversion, Vilnius, Lithuania) and operating at 1 kHz and a pulse energy of 0.2 mJ. About 120 µJ of the output is used to pump a tunable ORPHEUS optical parametric amplifier (Light Conversion, Vilnius, Lithuania) to generate pump pulses at 570 nm. About 20 µJ of the output is used to generate the second harmonic at 515 nm, which is then used to pump a sapphire substrate and generate white light continuum pulses in the 300-500 nm spectral region. The pump pulses are modulated using a mechanical chopper (Thorlabs, Newton, NJ) at 500 Hz, temporally delayed using a mechanical translation stage (PI, Auburn, MA) and focused on the sample at a slight angle to the probe pulses. The pump energy density is maintained between 10-100 µJ/cm^2^. The probe pulses are detected by an Acton spectrograph (Teledyne Princeton Instruments, Trenton, NJ) in combination with a high-speed spectroscopy camera (Stresing, Berlin, Germany). The transient absorption signal is then estimated by the analysis of the probe spectra measured synchronously with the optical chopper using a home-built LabVIEW routine.

### Direct measurement of RcoM-HBD-CCC O_2_ binding affinity (*K*_a,O2_)

RcoM-HBD-CCC bearing unliganded Fe(II) heme was generated by reacting chemically oxidized Fe(III) RcoM-HBD-CCC with excess sodium dithionite in a rigid glove box (Coy Lab Products, Grass Lake, MI) purged with a N_2(*g*)_ atmosphere and O_2_ levels maintained below 15 ppm. The chemically reduced protein sample was passed through a gravity size exclusion/desalting column pre-equilibrated with PBS. This column was pre-washed with 20 mM sodium dithionite to scavenge O_2_ in the column, followed by two rinses with PBS to remove excess sodium dithionite. Eluted RcoM-HBD-CCC bearing Fe(II) heme was loaded into a septum-sealed cuvette quartz cuvette (5 mL total volume) and transferred to a benchtop Cary 100 with sample holder maintained at 25 °C. Titration of the sample with O_2_ was accomplished by introducing air via gas-tight syringe into the cuvette headspace (2.2 mL) in 2 to 25 μL increments, inverting to mix, and allowing the mixture to equilibrate for 2 min prior recording UV-Vis spectral features. After introduction of 800 μL of air, no further spectral changes were observed. The cuvette was opened to air and allowed to react for 5 additional minutes before recording a final Fe(II)-O_2_ spectrum. Fractional O_2_-binding to Fe(II) RcoM-HBD-CCC heme was determined from changes in the absorbance at 561 nm. The partial pressure of O_2_ in the headspace was estimated assuming a pressure of 1 atm and atmospheric O_2_ concentration of 20.9%. A Henry’s Law constant of 1.73 μM•mmHg^-1^ was used to convert the equilibrium dissociation value *P*_50_ = 3.9 mmHg to *K*_d,CO_ = 6.7 μM (69).

### Determination of O_2_ binding dissociation rate constant *k*_off,O2_

The first-order rate constant for O_2_ dissociation from Fe(II) RcoM heme was determined by measuring replacement of O_2_ with CO using stopped-flow UV-Vis spectroscopy. Fe(II)-O_2_ RcoM-HBD-CCC (5 μM final heme concentration) in aerobic PBS were mixed in a 1:1 ratio in the stopped-flow mixing chamber and spectral features recorded for up to 10 s post-mixing. The change in Soret absorbance at 423 nm was fit to single-exponential decay functions to determine observed pseudo-first order rate constant, and the final rate constant value was reported as the average of fitted rate constants from at least four traces.

### CO scavenging from red blood cell-encapsulated HbCO *ex vivo*

CO transfer from RBC-encapsulated HbCO to cell-free RcoM-HBD-CCC was carried out as described previously for other CO scavenger (21, 22). Blood was collected from healthy C57BL/6J mice and utilized for *ex vivo* CO transfer experiments on the same day. RBCs were separated from plasma by centrifugation a 2,000 × *g* for 1 min at 4 °C, followed by three successive washes with ice cold PBS, followed by centrifugation at 2,000 × *g* for 1 min. To generate CO-saturated (>95%) Hb, washed red cells were diluted five-fold into CO-saturated PBS with 10 mM sodium dithionite and allowed to react for 5 min on ice. After washing cells to remove excess CO, we mixed CO-saturated RBCs with active, oxyferrous RcoM-HBD-CCC and incubated for 30 min. At prescribed time intervals, we removed and centrifuged aliquots of the suspension to separate RBCs from RcoM-HBD-CCC and thereby halt CO transfer. After collecting aliquots, we recorded UV-Vis spectra of the supernatant and lysed RBCs. Using basis spectra for CO-bound and CO-free Hb and RcoM-HBD-CCC, we quantified the relative percentage of CO bound to each hemoprotein by spectral deconvolution.

### Preparation of Fe(II)-O_2_ StHb

2,3-diphosphoglycerate stripped hemoglobin (StHb) bearing >99% Fe(II)-O_2_ heme was isolated from expired, leukoreduced human donor packed RBC units as previously reported (21, 22).

### *In vivo* CO scavenging during acute inhaled CO poisoning

All animal protocols were reviewed and approved by the Institutional Animal Care and Use Committees at the University of Pittsburgh and University of Maryland Baltimore. Mice were housed at the University of Pittsburgh Division of Laboratory Animal Resources or the University of Maryland Baltimore Veterinary Resources in rooms maintaining 12/12 h light/dark cycles at 20–26 °C and 30–70% relative humidity as prescribed by the National Research Council.

A murine model of severe CO poisoning was adapted from prior studies assessing CO scavenging efficacy during acute CO poisoning (21, 22). To facilitate comparison to prior studies, all animal experiments were conducted using male C57BL/6J mice (Jackson Laboratories), aged between 12-14 weeks. Mice were anesthetized with 1%–3% isoflurane throughout all surgical procedures. After making a single incision on the neck, a tracheal tube was introduced via a tracheotomy for mechanical ventilation with a 20-gauge cannula. Catheters (0.025” × 0.012”) were inserted into the right carotid artery and jugular vein, and the neck incision closed with 6-0 sutures. After surgical instrumentation, anesthetized mice were moved into a fume hood for the remainder of the procedure to minimize CO exposure for researchers. Body temperature, monitored by rectal thermometer, was maintained by at 37 °C throughout the procedures using a heated platform connected to a water pump (Adroit Medical Systems, Loudon, TN).

Volume-controlled mechanical ventilation of air with 1.5% isoflurane (fraction of inspired oxygen 21%, tidal volume 8.8 mL/kg body weight, 175 breaths per minute) was applied with a small animal ventilator (MiniVent, Type 845; Hugo Sachs). Arterial blood pressure was continuously recorded (PowerLab 8/35, ADInstrument) via the arterial catheter every 1 ms. The MAP and pulse rate were calculated by Labchart software (ADInstrument, v7). To minimize experimental noise, MAP values were plotted at 10 s intervals with each datapoint representing the average of 10 values. Animals were monitored for stable heart rate and blood pressure for at least 20 min of ventilation prior to CO exposure. After the stabilization period, three percent CO in air was administered for 1.5 min, followed by continued ventilation with air. At prescribed time intervals, 15 μL blood samples were drawn via arterial catheter and processed as described below.

RcoM-HBD-CCC bearing Fe(II)-O_2_ heme (or PBS vehicle) was delivered via jugular venous catheter over an infusion period of 2 min at an infusion volume of 10 mL/kg body weight. Infusion was initiated 4.5 minutes after initial CO exposure. RcoM-HBD-CCC (>99% Fe(II)-O_2_) was infused at Fe(II)-O_2_ heme concentrations of 1.5 mM, 3 mM, and 6 mM, corresponding to total doses of 255 mg/kg, 510 mg/kg, and 1020 mg/kg, respectively. StHb (>99% Fe(II)-O_2_) was infused at a 8.5 mM dose, corresponding to at total dose of 850 mg/kg.

### Quantification of hemoprotein concentrations and species distribution

Mouse blood samples (15 μL) drawn in CO poisoning studies were immediately centrifuged at 10,000 ×*g* for 1 minute to separate RBCs from plasma. RBC pellets were resuspended with PBS 1ml and centrifuged again at the same speed for 1 minute, followed by removal of supernatant to avoid plasma residue. The plasma and RBC pellet samples were flash frozen separately on dry ice and stored at –80°C. Analysis of Hb and RcoM species were carried out separately as centrifugation allowed for complete separation of RBC-encapsulated Hb and RcoM-HBD-CCC in plasma. To determine the fraction of CO-bound hemoglobin, each frozen RBC pellet was thawed and lysed by dilution into a hypotonic 20 mM sodium dithionite solution. RBC lysates were immediately analyzed by UV-Vis spectroscopy. Heme speciation for extracellular RcoM-HBD-CCC was analyzed by dilution of corresponding plasma samples in PBS and recording spectral features, then adding excess sodium dithionite and re-recording absorption spectra. All spectra were recorded from 450–700 nm. Spectral deconvolution was employed to determine the concentrations and relative fractions of hemoprotein species for RBC-derived Hb and plasma-derived RcoM-HBD-CCC using standard references for Fe(III), Fe(II), Fe(II)-O_2_, and Fe(II)-CO species.

### Hemodynamic analysis of hemoprotein infusion in non-CO poisoned mice

The exact same surgical procedure utilized in the above CO poisoning model above was employed to compare hemodynamic effects of hemoprotein infusion between RcoM-HBD-CCC and StHb, with the exception that room air was supplied during mechanical ventilation for the entire experiment without exposure to CO_(*g*)_. RcoM-HBD-CCC and StHb (>99% Fe(II)-O_2_) were infused at a concentration of dose of 800 mg/kg (50 μmol/kg heme; *n*=3) and of 1020 mg/kg protein (60 μmol/kg heme; *n*=4), respectively.

### Estimation of *k*_NOD,RcoM_ through competitive reaction with hemoglobin

In a cuvette under aerobic conditions, 30 μM Fe(II)-O_2_ RcoM-HBD-CCC was mixed with 30 μM Fe(II)-O_2_ StHb at 20 °C in PBS. A stock ProliNONOate solution (1 mM in 10 mM NaOH) was added to a final concentration of 30.2 μM NO, the solution inverted several times to mix, and mixture allowed to react for 5 min. The reaction progress was monitored every minute until no further spectroscopic changes were observed, approximately 5 minutes.

### RcoM-HBD-CCC safety and tolerability *in vivo*

Healthy (non-CO-poisoned) C57BL/6J mice were anesthetized with inhaled isoflurane (2-3%) and body temperature maintained at 37°C. Fe(II)-O_2_ RcoM-HBD-CCC in 10 mM PBS, pH 7.4 was infused via lateral tail vein catheter over 30 minutes by an injection pump. The catheter was removed, followed by ligation of the vessel and closure of the incision. Mice were then placed in cages and observed for 48 hours, monitoring activity, nesting, and daily weights. The mice were euthanized after 48 hours. At necropsy, blood samples were sent for serum chemistry measurement (Marshfield Labs, Cleveland, OH), and organs were harvested and stored at −80 C for subsequent analysis of RcoM accumulation by Western Blot.

* Concentrations based on holoprotein concentrations derived from heme quantitation using the pyridine hemochromagen assay

## AUTHOR CONTRIBUTIONS

M.R.D. conceptualized the scavenger, designed and performed experiments, analyzed data, and wrote the initial and final drafts of the paper. A.W.D., Q.X., X.C., A.G., J.H., and J.J.R. K.A.B., performed key experiments and/or analyzed data. E.A, K.B.U., S.R.B, A.R.S.K., A.B., and D.B.K.-S. designed, carried out, and interpreted results from picosecond spectroscopy experiments. All authors reviewed and edited the manuscript. M.T.G. and J.T. directed the research, designed experiments, interpreted data, and share senior authorship.

## ACKNOWLEDGEMENTS

This work is supported by the NIH (grants T32HL110849, F32HL162381, and K99HL168224 to MRD; grant K08 HL136857 to JJR; grant R01 HL125886 to MTG and JT; and grant R42 ES031993 to JT and JJR through collaboration with Globin Solutions Inc.), the DOD (grant W81XWH2210198 to JJR and JT), the Martin Family Foundation to MTG, and the Parker B. Francis Foundation to JJR.

