## Supplemental Materials for "Engineering a highly selective, hemoprotein-based scavenger as a carbon monoxide poisoning antidote with no hypertensive effect"

* corresponding authors:

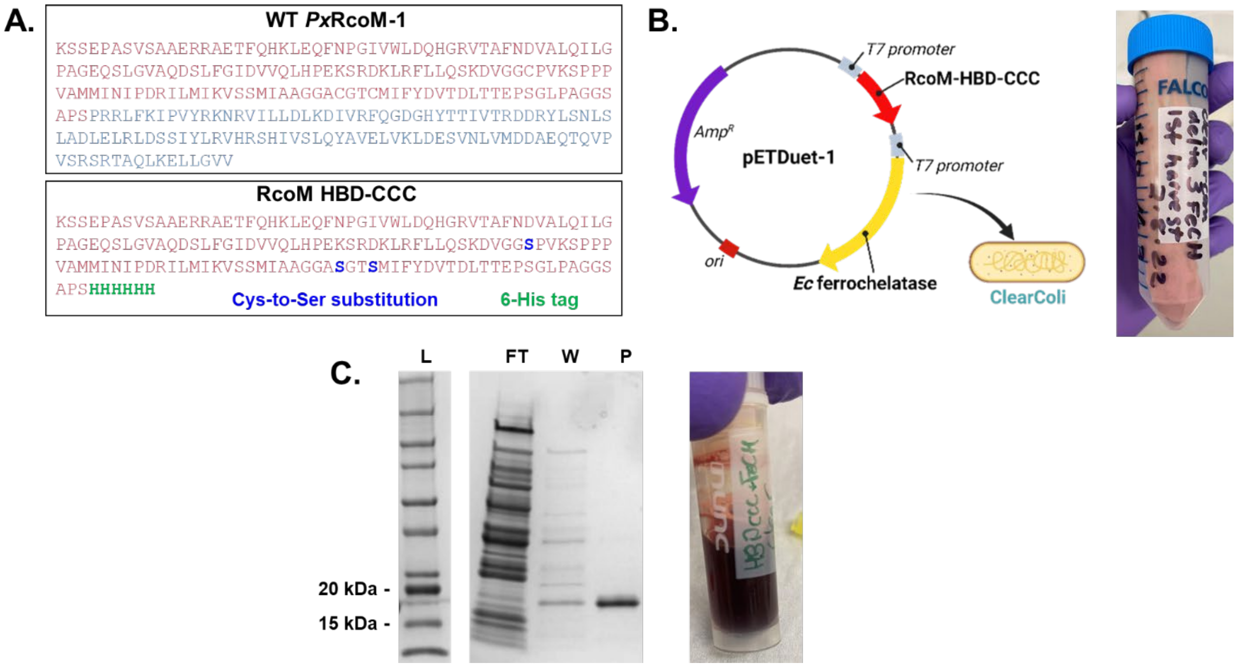


**Figure S1. (A)** Amino acid sequence for WT RcoM-1 from *Paraburkholderia xenovorans* (*Px*RcoM-1) and the RcoM truncate variant, RcoM HBD-CCC. **(B)** CO-expression vector bearing open reading frames for RcoM HBD-CCC and *E. coli* ferrochelatase (left) and representative image of harvested E. coli cell paste following overexpression using this vector in ClearColi cells (left). **(C)** Representative SDS-PAGE of RcoM HBD-CCC (MW=17.0 kDa) fractions after Ni-NTA column chromatography: flow through (**FT**), wash (**W**), and purified (**P**) protein. **(D)** Representative images of bacterial cell paste after overexpression of RcoM HBD-CCC (left) and purified protein sample (right).

**Table S1.** Representative expression parameters for RcoM HBD-CCC using different expression vectors and *E. coli* strains.

| Expression plasmid, strain | Trial | wet cell mass (g/L culture) | $[\text{protein}]_{total}$ (mM) | [holoprotein] (mM) | Heme loading (%) | holoprotein yield (mg/L culture) |
|---|---|---|---|---|---|---|
| pET28a (RcoM HBD-CCC), soluBL21 | 1 | 11.0 | 4.84 | 0.99 | 20 | 43.7 |
| | 2 | 10.4 | 5.3 | 0.895 | 17 | 46.5 |
| | 3 | 15.0 | 8.37 | 1.32 | 16 | 49.2 |
| | **Average** | **12.1** | **6.2** | **1.1** | **18** | **46.5** |
| pETDuet (RcoM HBD-CCC + *Ec*FeCH), soluble21 | 1 | 7.5 | 2.88 | 3.55 | 123 | 15.4 |
| | 2 | 10.4 | 2.36 | 3.2 | 136 | 11.1 |
| | 3 | 10.7 | 6.05 | 6.54 | 108 | 4.2 |
| | 4 | 6.5 | 3.74 | 3.81 | 102 | 5.0 |
| | 5 | 5.4 | 2.59 | 2.73 | 105 | 7.1 |
| | **Average** | **8.1** | **3.5** | **4.0** | **115** | **8.6** |
| pETDuet (RcoM HBD-CCC + *Ec*FeCH), ClearColi | 1 | 3.0 | 2.84 | 2.46 | 87 | 1.2 |
| | 2 | 2.8 | 1.92 | 2.031 | 106 | 1.0 |
| | 3 | 2.4 | 2.77 | 3.707 | 134 | 2.4 |
| | 4 | 2.8 | 1.27 | 1.236 | 97 | 1.3 |
| | 5 | 2.4 | 3.19 | 3.32 | 104 | 3.1 |
| | **Average** | **2.7** | **2.4** | **2.6** | **106** | **1.8** |

**Table S2.** Summary of spectroscopic features for RcoM HBD-CCC heme in PBS at 25 °C.

| RcoM species | Soret peak, nm | β band, nm<br>ε, $mM^{-1} cm^{-1}$ | α band, nm<br>ε, $mM^{-1} cm^{-1}$ |
|---|---|---|---|
| Fe(III) | 415 | 537<br>(10.6) | 565<br>(8.8) |
| Fe(II) | 424 | 531<br>(12.5) | 562<br>(19.9) |
| Fe(II)-$O_2$ | 422 | 540<br>(14.8) | 573<br>(14.7) |
| Fe(II)-CO | 422 | 541<br>(13.6) | 572<br>(12.6) |
| Fe(II)-NO | 421 | 546<br>(12.4) | 579<br>(12.4) |

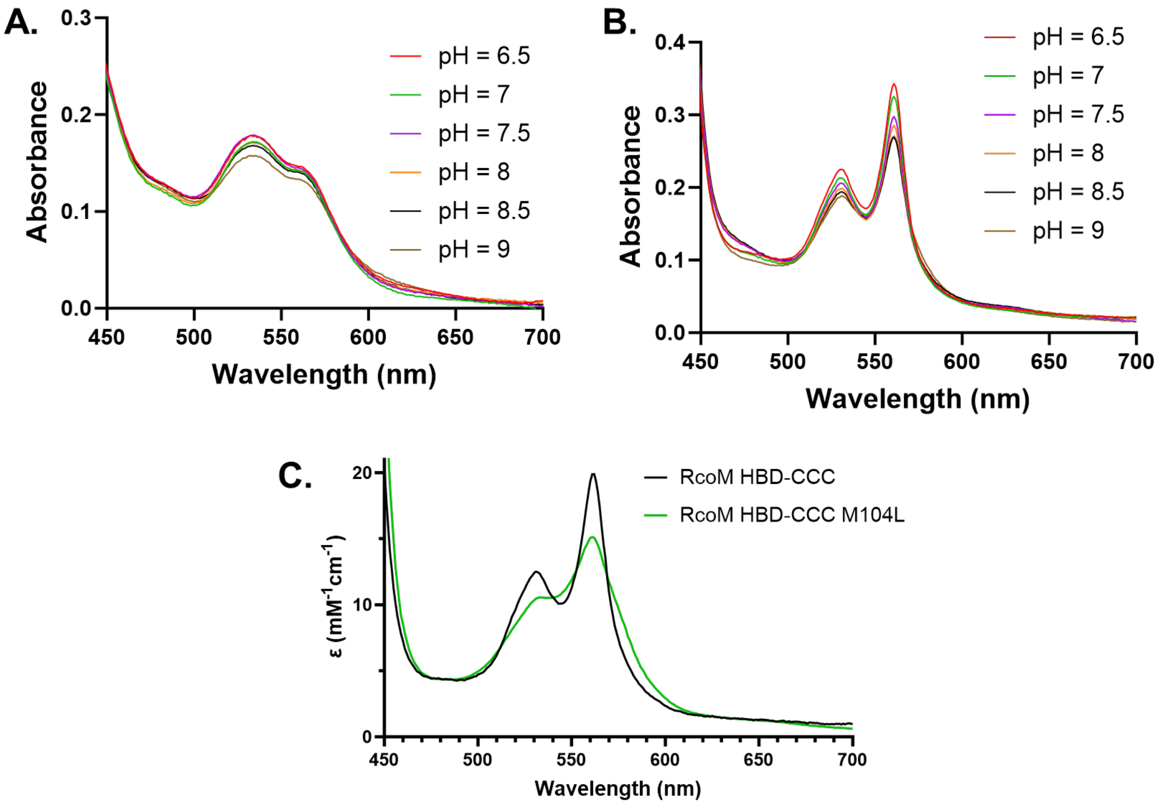


**Figure S2.** Assessment of pH-dependent changes in visible spectral features for Fe(III) RcoM HBD-CCC **(A)** and Fe(II) RcoM HBD-CCC **(B)**. Comparison of spectroscopic features for unliganded, Fe(II) RcoM HBD-CCC and RcoM HBD-CCC M104L variant **(C)**. Spectra were recorded at 25 °C in 50 mM buffer, 150 mM NaCl where buffer was MES for pH = 6.5; MOPS for pH = 7.0 and 7.5, and TRIS for pH = 8.0, 8.5, and 9.0.

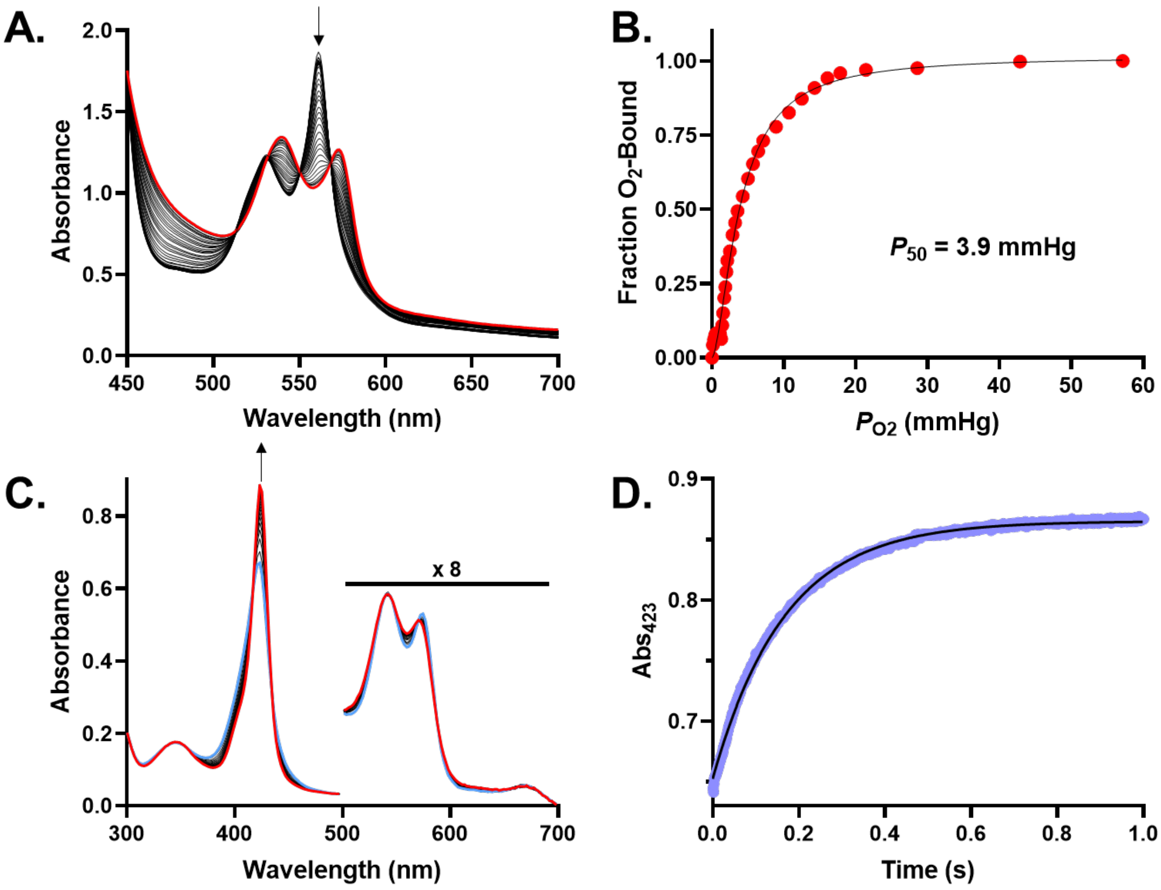


**Figure S3.** Oxygen binding properties of RcoM-HBD-CCC in PBS at 25 °C. **(A)** Thermodynamic oxygen binding of Fe(II) RcoM-HBD-CCC (20 μM), as measured by changes in the visible (Q-band) region. **(B)** Oxygen binding curve for RcoM-HBD-CCC, determined from spectral deconvolution of visible absorbance features. **(C)** Oxygen dissociation kinetics were measured by rapid mixing of Fe(II)-$O_2$ RcoM-HBD-CCC with CO-saturated buffer in a stopped-flow apparatus. **(D)** The change in Soret absorbance at 423 nm was fit to a single-exponential curve to yield a first-order rate constant of 5.9 $s^{-1}$.

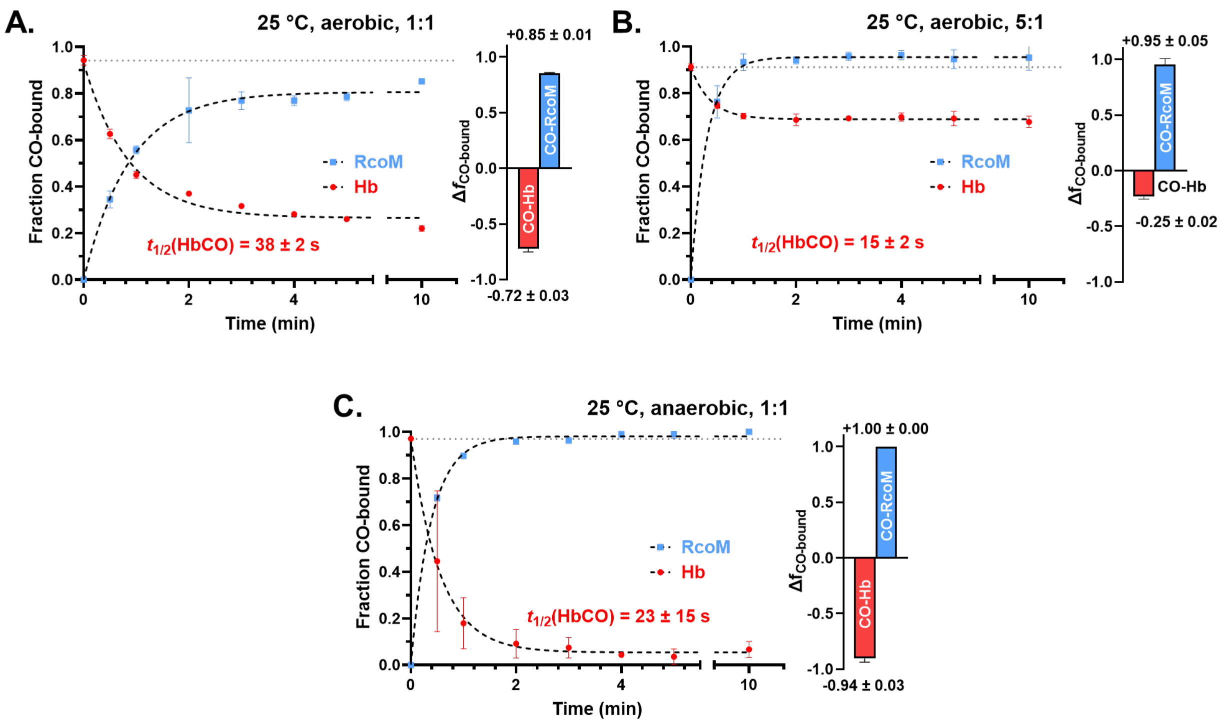


**Figure S4.** *In vitro* CO scavenging in PBS (pH 7.4) at 25 °C under **(A)** aerobic conditions (100 μM RcoM and 100 μM COHb), **(B)** anaerobic conditions (130 μM RcoM and 130 μM COHb), and **(C)** aerobic conditions with 5-fold excess COHb (1 mM) relative to RcoM (200 uM).

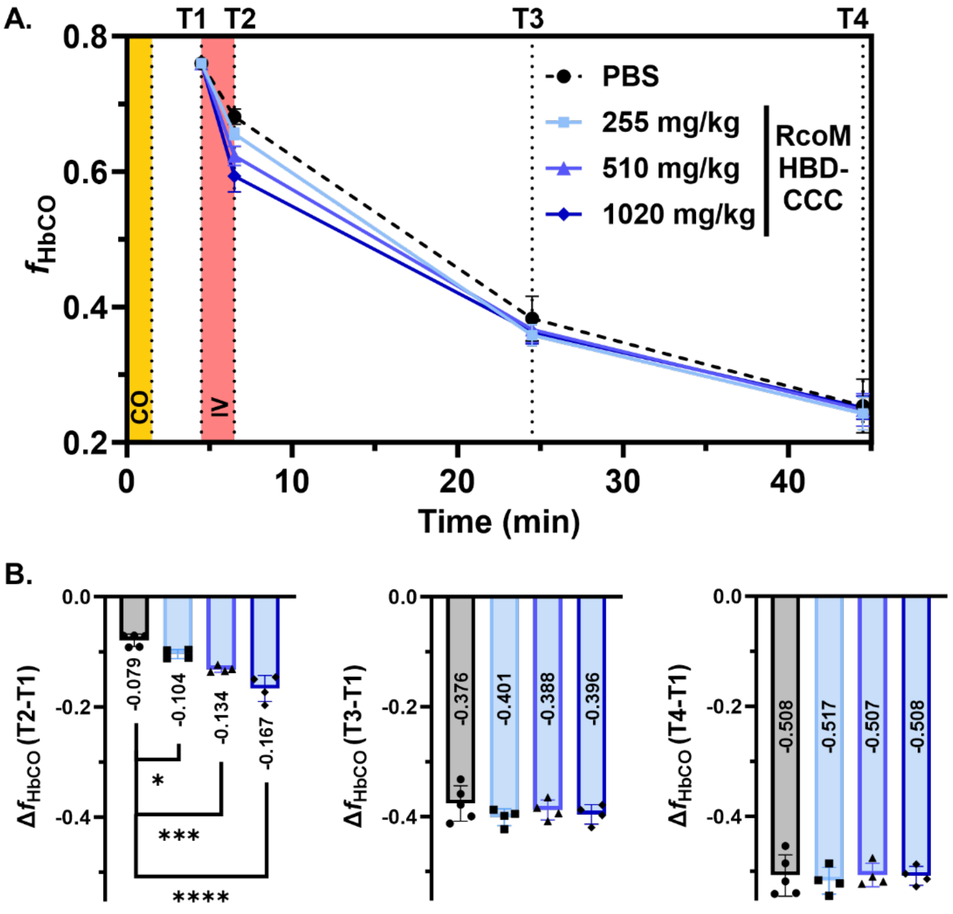


**Figure S5. (A)** Changes in the fraction of circulating HbCO ($f_{HbCO}$) after CO exposure and delivery of RcoM-HBD-CCC scavenger. All $f_{HbCO}$ values were adjusted to a starting value of 0.76, the average initial $f_{HbCO}$ at T1 across all treatment groups. **(B)** Comparison of CO clearance, $\Delta f_{HbCO}$ , at different time points throughout the severe CO poisoning model. Statistical significance between vehicle and treatment groups was assessed using ordinary one-way ANOVA (*, $p<0.05$; ***, $p<0.001$; ****, $p<0.0001$).

**Table S3.** Summary of hemodynamic data for severe, nonlethal CO poisoning model.

| | | Average MAP, mmHg @ (SD, mmHg) | | | | ΔMAP, mmHg @ (SD, mmHg)** | | | | | |
|---|---|---|---|---|---|---|---|---|---|---|---|
| | **treatment** | ***t = 0**** | ***min (pre-infusion)*** | ***min (post-infusion)*** | ***max (post-infusion)*** | ***t* = 10 min** | ***t* = 20 min** | ***t* = 30 min** | ***t* = 40 min** | **min** | **max** |
| | **PBS** | 84.5 (4.3) | 50.0 (6.8) | 82.9 (6.2) | 103.9 (8.0) | 4.7 (1.2) | 6.9 (1.3) | 5.8 (2.1) | 4.7 (2.5) | -1.2 (3.5) | 17.1 (5.0) |
| **RcoM-HBD-CCC** | **15 μmol/kg heme** | 83.4 (5.4) | 48.2 (3.8) | 83.6 (3.0) | 94.0 (3.8) | 3.8 (6.9) | 7.3 (4.8) | 7.0 (4.0) | 5.2 (4.4) | 0.7 (5.3) | 11.1 (5.4) |
| | **30 μmol/kg heme** | 81.5 (1.5) | 49.1 (2.7) | 76.8 (1.5) | 87.2 (1.1) | 0.2 (1.6) | 4.1 (1.8) | 2.9 (1.8) | 2.1 (1.3) | -4.4 (3.0) | 6.0 (1.2) |
| | **60 μmol/kg heme** | 85.1 (5.4) | 46.6 (3.6) | 80.3 (2.4) | 93.0 (4.9) | 1.4 (5.9) | 1.7 (7.5) | 1.5 (6.7) | 1.0 (6.4) | -4.3 (5.4) | 8.4 (6.1) |
| **StHb** | **50 μmol/kg heme** | 82.8 (1.7) | 39.9 (10.7) | 79.9 (9.7) | 105.9 (8.4) | 8.8 (6.6) | 14.7 (5.7) | 19.4 (9.2) | 16.7 (8.0) | -2.9 (8.2) | 23.1 (8.2) |

*mean arterial pressure (MAP) for each biological replicate was averaged over the first 30 s of blood pressure monitoring. The summarized data represent an average across all biological replicates in each treatment group at *t* = 0.

**$\Delta MAP = MAP_{tx} - MAP_{t0}$ where x = 10, 20, 30, or 40 min after the start of blood pressure monitoring, the global minimum blood pressure following infusion (min), or the global maximum blood pressure following infusion (max). For each biological replicate, the MAP value at each time point was determined by averaging blood pressure over a one-minute window centered on that time point. The summarized data represent an average across all biological replicates in each treatment group at each time point.

**Table S4.** Comparison of hemodynamic data for infusion of hemoprotein-based CO scavengers in a non-CO poisoning murine model.

| | Average MAP, mmHg @ (SD, mmHg) | | | | ΔMAP, mmHg @ (SD, mmHg)** | | | | | |
|---|---|---|---|---|---|---|---|---|---|---|
| treatment | *t = 0** | *min (pre-infusion)* | *min (post-infusion)* | *max (post-infusion)* | *t* = 10 min | *t* = 20 min | *t* = 30 min | *t* = 40 min | min | max |
| **PBS** | 86.4 (2.8) | 83.9 (3.0) | 76.9 (3.4) | 95.9 (6.8) | -2.1 (4.5) | -1.1 (2.0) | 0.3 (1.4) | 2.8 (0.1) | -9.5 (6.8) | 9.5 (5.1) |
| **RcoM 30 μmol/kg heme** | 85.8 (5.2) | 83.6 (6.0) | 73.6 (8.1) | 87.3 (3.6) | -5.5 (1.3) | -2.8 (2.8) | -2.3 (2.4) | -0.5 (2.0) | -12.2 (6.2) | 1.5 (1.7) |
| **StHb 30 μmol/kg heme** | 85.9 (3.6) | 84.5 (4.0) | 88.4 (2.7) | 98.8 (2.2) | 8.0 (4.3) | 6.3 (5.6) | 9.5 (7.3) | 6.6 (8.7) | 2.7 (6.3) | 13.1 (5.9) |

*mean arterial pressure (MAP) for each biological replicate was averaged over the first 30 s of blood pressure monitoring. The summarized data represent an average across all biological replicates in each treatment group at $t = 0$.

**$\Delta MAP = MAP_{tx} - MAP_{t0}$ where x = 10, 20, 30, or 40 min after the start of blood pressure monitoring, the global minimum blood pressure following infusion (min), or the global maximum blood pressure following infusion (max). For each biological replicate, the MAP value at each time point was determined by averaging blood pressure over a one-minute window centered on that time point. The summarized data represent an average across all biological replicates in each treatment group at each time point.

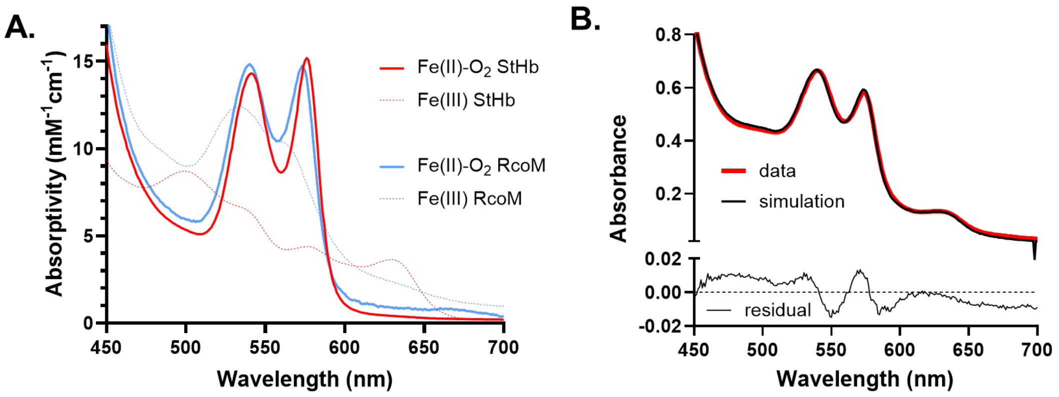


**Figure S6.** Determination of $k_{NOD,RcoM}$ by reaction competition with StHb . **(A)** Comparison of spectroscopic data for Fe(III) RcoM-HBD-CCC (blue, dashed line), Fe(II)-$O_2$ RcoM-HBD-CCC (blue, solid line), Fe(III) StHb (red, dashed line), Fe(II)-$O_2$ StHb (red, solid line). **(B)** Representative spectral deconvolution data (black line) in which the relative contributions of the four reference spectra from panel A were fit using experimental NO scavenging data (red line).

**Table S5.** Summary of deconvolution data for NO dioxygenation competition experiment after 5 minutes of reaction between ~30 μM Fe(II)-$O_2$ RcoM, 30 μM Fe(II)-$O_2$ StHb, and 30 μM NO..

| trial | [oxy RcoM] (μM) | [oxy Hb] (μM) | [Fe(III) RcoM] (μM) | [Fe(III) Hb] (μM) | $[RcoM]_{tot}$ (μM) | $[StHb]_{tot}$ (μM) | total NO |
|---|---|---|---|---|---|---|---|
| 1 | 25.7 | 6.3 | 4.5 | 27.1 | 30.2 | 33.4 | 31.6 |
| 2 | 25.9 | 3.5 | 4.5 | 27.8 | 30.4 | 31.3 | 32.3 |
| 3 | 25.7 | 5.4 | 4.6 | 27.8 | 30.3 | 33.2 | 32.4 |
| 4 | 25 | 4.6 | 5.1 | 26.2 | 30.1 | 30.8 | 31.3 |

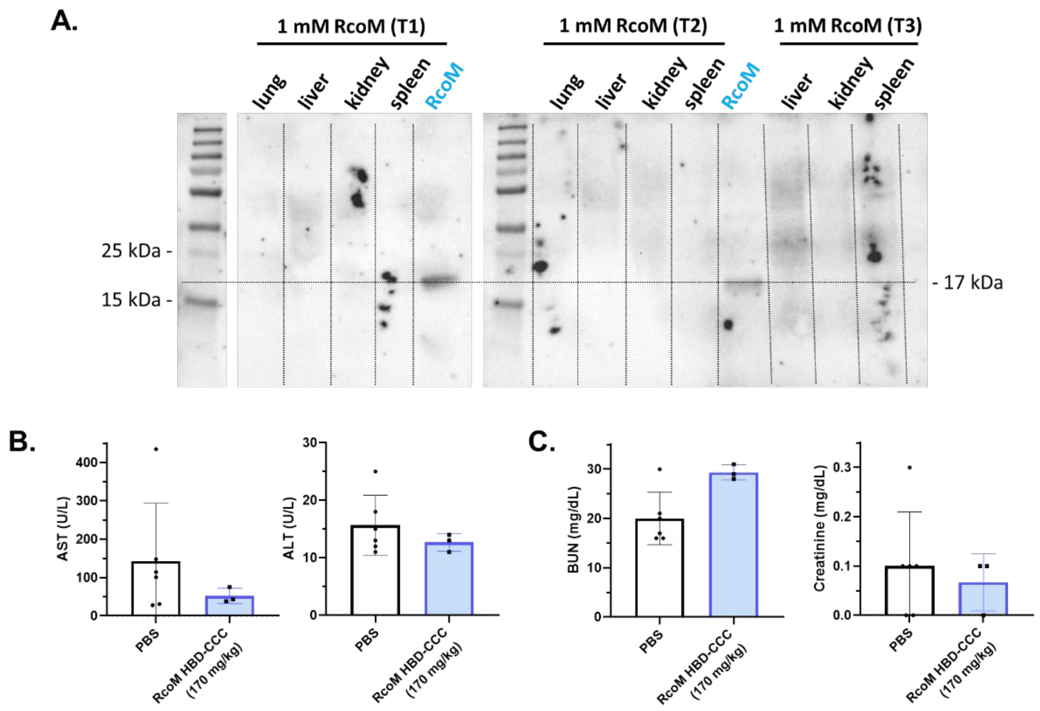


**Figure S7.** RcoM-HBD-CCC safety and tolerability. (A) Western blots show lack of accumulation of scavenger in tissue homogenates from non-CO poisoned animals administered RcoM-HBD-CCC (170 mg/kg protein, 1.5 μmol/kg heme) in the 48-hour toxicity screening model. RcoM-HBD-CCC was detected using anti-6xHis antibodies. Positive control lanes contain heterogously expressed RcoM-HBD-CCC purified to homogeneity as described in the Materials and Methods section. Quantification of plasma biomarkers indicative of liver (B) and kidney (C) function for animals administered RcoM-HBD-CCC ($N$=3) and PBS ($N$=6) vehicle in the 48-hour toxicity screening model. No significant differences were observed between treatment groups, as assessed using one-way analysis of variance (ANOVA) testing.

**Table S6.** Primers for Gibson assembly cloning and site-directed mutagenesis.

| | |
|---|---|
| **Gibson assembly primers** | |
| RcoM_Gib_fw | ttaagaaggagatataccatgaagtctagcgagcctg |
| RcoM_Gib_rv | atggctgctgcccattcaatggtggtggtggtg |
| pET28a_Gib_fw | atgggcagcagccatcatc |
| pET28a_Gib_fw | ggtatatctccttcttaaagttaaacaaaattatttctagag |
| **Site-directed mutagenesis primers** | |
| C127S_fw | TGGCGGCGCGAGTGGCACCTGTA |
| C127S_rv | GCAGCAATCATGCTCGATACCTTAATCATCAGGATG |
| C130S_fw | GTGTGGCACCAGTATGATTTTCTATGATGTAACAGAC |
| C130S_rv | GCGCCGCCAGCAGCAATC |
| M104L_fw | GCCGGTGGCGCTGATGATCAACATTC |
| M104L_rv | GGCGGGCTTTTAACCGGG |
